# Decoding the avian missing gene mystery: dot chromosomes unmask extensive gene loss and novel genetic instability

**DOI:** 10.1101/2025.09.16.676548

**Authors:** Tomáš Hron, Dalibor Miklík, Jan Pačes, Petr Pajer, Vladimír Pečenka, Jiří Hejnar, Jiří Nehyba, Daniel Elleder

**Affiliations:** Institute of Molecular Genetics of the Czech Academy of Sciences, Vídeňská 1083, 14220 Prague, Czech Republic

## Abstract

The apparent absence of numerous conserved vertebrate genes from avian genomes has puzzled researchers for over a decade. In recent years, a subset of these genes has been identified; however, their sequences are unusually problematic, often evading detection by standard sequencing technologies. This limitation has hindered detailed investigation of the phenomenon—until recent progress in long read technologies, which are more robust against sequencing biases.

This enabled us to classify real gene losses extensively, which strikingly revealed that a large number of the genes residing on so-called dot chromosomes were indeed lost during avian evolution. We demonstrate that dot microchromosomes—small, repeat-dense avian chromosomes—harbor widespread gene attrition, with 29% of ohnologs (duplicates from ancestral genome doublings) eliminated, far exceeding rates on other chromosomes. Moreover, we reveal that genes retained on these dot chromosomes exhibit a previously undescribed form of dynamic genetic instability. This instability, which we term sequence stuttering, is characterized by a massive expansion of short sequences within intronic regions. Intriguingly, in some cases, the expanding sequences appear to originate from neighboring exons. As a result, intron lengths vary extensively among individual chickens, suggesting that these events are evolutionarily recent.

Since this phenomenon has not been reported in any other vertebrate species, our findings lay the groundwork for future research into its underlying mechanisms, evolutionary implications, and potential identification of similar loci across vertebrate genomes.

**SIGNIFICANCE:** This study reveals extensive gene loss and a novel genetic instability, termed “sequence stuttering,” on avian dot microchromosomes, providing new insights into avian genomic evolution. Approximately 29% of ohnologs on dot chromosomes are absent compared to other vertebrate chromosomes, possibly due to elevated GC content and recombination rates in euchromatin regions. This significant gene loss highlights dot chromosomes as hotspots for genomic reduction, potentially shaping avian-specific traits. Additionally, sequence stuttering—characterized by extensive intronic repeat expansions, sometimes incorporating exonic sequences—introduces marked length polymorphism within chicken populations, suggesting ongoing evolutionary dynamics. These findings underscore the unique role of dot chromosomes in avian genome evolution, emphasizing their contribution to genetic diversity and adaptation. This work lays the foundation for further investigation into the molecular mechanisms driving these phenomena and their broader implications for vertebrate genomic evolution.

## INTRODUCTION

Even before the first release of the chicken genome assembly it was known that avian genes have on average higher guanine and cytosine (GC) content than mammalian genes (International Chicken Genome Sequencing Consortium 2004). The avian karyotype was also known to be quite distinct compared to other vertebrates by containing a large number of small GC-rich chromosomes, known as microchromosomes (Smith et al. 2022; Burt 2002). In 2015, we reported that a subset of avian genes has extremely high GC content, often concentrated in short GC-rich sequence stretches (Hron et al. 2015). These sequences are extremely difficult to amplify by PCR and process by next generation sequencing (NGS). Therefore, these genes were often falsely considered missing, and we named them “hidden avian genes”. Since then, analysis of deep-coverage sequencing data, optimized PCR amplification methods, and other dedicated methods led to the identification of other avian genes in this category, including, for example, avian tumor necrosis factor alpha (TNF-α), erythropoietin (EPO), leptin, LAT, FOXP3, and interferon regulatory factors (IRF3, IRF9) (Rohde et al. 2018; Janusova et al. 2023; Burkhardt et al. 2022; Ungrová et al. 2025; “Identification of a GC-Rich Leptin Gene in Chicken” 2016; Hron et al. 2015). Systematic analyses of transcriptomic and genomic data enabled large scale identification of multiple additional avian “hidden” genes (M. Li et al. 2022; Laine et al. 2019; Zhu et al. 2023; Hu et al. 2024; Bornelöv et al. 2017; Huttener et al. 2021; Zhao, Yin, and Hou 2025).

A definitive explanation for the GC-rich and atypical sequence characteristics of some avian genes has not been provided. It is considered to be partially caused by the specifics of avian GC-biased gene conversion (gBC) and is known to be more pronounced on microchromosomes (Botero-Castro et al. 2017; Huttener et al. 2021). The original hypothesis of large-scale gene loss in the avian genome has become more controversial (Lovell et al. 2014). Overall, the gene loss was considered to be mainly a technical problem of gene identification, and most missing avian genes were expected to be discovered (Lovell et al. 2014; M. Li et al. 2022; Zhao, Yin, and Hou 2025).

In 2021, the reference RefSeq annotated chicken genome GRCg7b was released (Smith et al. 2022). Despite a high quality of assembly in standard chromosomes and standard gene annotations, this genome does not contain a significant portion of highly repetitive, usually GC-rich regions, including one entire microchromosome and over 40% of the assembled sequences of 18 other small microchromosomes. In the last couple of years, large progress was enabled by improvements in nanopore sequencing, which has very low GC bias and long reads (Browne et al. 2020). Importantly, a new high quality complete (telomere-to-telomere) chicken genome was published (Huang, Xu, Bai, et al. 2023). Only provisional XenoRefSeq mRNA gene annotation, built on the blat alignments of GenBank vertebrate and invertebrate reference mRNAs, is provided as a track in the UCSC genome browser database for this genome (Perez et al. 2025). Although this annotation is inaccurate in the exact positions of individual exons, it represents the most complete chicken gene annotation to date. Importantly, the 2023 genome analysis introduced a special category of microchromosomes called dot chromosomes, on the basis of epigenetic modifications and genomic repeat content (Huang, Xu, Bai, et al. 2023).

Ten dot chromosomes are an integral part of the chicken karyotype of 39 chromosome pairs. The karyotype is composed of macrochromosomes that, depending on classification, typically include the 6-10 largest chromosomes and of the remaining smaller chromosomes classified as microchromosomes (Smith et al. 2022). Newly defined dot microchromosomes, constituted by chromosomes 16, 29-32, and 34-38, are by their size at the bottom of the microchromosome size scale and are distinguishable from other microchromosomes by very high repeat content, increased DNA methylation, and a specific histone methylation pattern. Further, they are strictly compartmentalized into euchromatin, centromere-distal heterochromatin, and centromere-proximal heterochromatin. Protein-coding genes are located in euchromatin and centromere-distal heterochromatin. Euchromatin genes are highly expressed one-copy-per-chromosome genes, while heterochromatin-localized genes have lower expression and are represented by a few classes of tandemly duplicated genes. Euchromatin specifics further include a high content of tandem simple repeats, depletion of interspersed repeats, variable DNA methylation level, and a variable but, on average, high GC content. In contrast, the centromere-distal heterochromatin has lower content of tandem simple repeats but contains numerous interspersed repeats, has less variable but, on average, higher DNA methylation, and somewhat lower average GC content (Huang, Xu, Bai, et al. 2023).

Dot chromosomes were so far defined only in chicken. However, the high conservation of avian karyotypes, both in chromosome number and chromosome gene content, suggests that dot-like chromosomes are present in the majority of avian species. Fifty percent of birds, including the chicken, have from 39 to 41 chromosome pairs, with the most common number being 40 (Degrandi et al. 2020). Individual avian chromosomes share synteny across multiple avian species (Waters et al. 2021). The presence of chromosomes with chicken dot chromosome-like gene content can be ascertained in annotations of high quality genomes in the current NCBI database including the genomes of ostrich, rhea, kiwi, nicobar pigeon, oriental stork, american crane, common cuckoo, and zebra finch.

In this study, the complete nature of the current chicken genome assembly as well as the availability of high-quality genomes of other avian species enabled us to classify real gene losses for the first time, catalogue them extensively, and map them to dot chromosomes. Additionally we described a so far unreported form of dynamic genetic instability on dot chromosomes in the chicken, which might be connected to the massive decay of avian genes in specific regions of their genomes.

## RESULTS

### Dot chromosomes as hotspots of gene loss in avian genomes

To explore the extent of evolutionary loss of avian genes and determine the principal chromosomal locations of these losses, we took advantage of the compartmentalization of gene content of vertebrate genomes into syntenic gene clusters, paralogons. These syntenic groups originated by two whole genome duplications (WGD) in ancestors of modern jawed vertebrates. Despite extensive gene loss after each WGD, many of the double-duplicated genes survived and exist today in vertebrate genomes as two, three, or four paralogs. These surviving paralogs are called ohnologs and the analysis of their synteny in different vertebrate species led to the definition of 68 paralogons that represent quadrupled 17 chromosomes of the putative vertebrate ancestor (Simakov et al. 2020; Lamb 2021).

To investigate gene losses in birds, the presence of ohnologs was quantified in the genomes of several taxonomic groups of tetrapod vertebrates and compared with the presence of ohnologs in avian species. Focusing on ohnologs had several advantages. Ohnologs are genes with a determined “date of birth” because they all originated in the last whole-scale duplication event, the second WGD common to all jawed vertebrates. Because the second WGD was an allopolyploidy, we understand the WGD event starting at the time of speciation that separated two vertebrate lineages that later hybridized. Also, the occurrence of ohnologs could be compared in pairs of syntenic clusters where each pair arose by the second WGD from genes located on a single chromosome. We will refer to those paralogons in the text that follows as ‘isoparalogons’. Finally, the avian karyotype is highly conserved and has close similarities with the karyotype of the hypothetical jawed vertebrate ancestor (Waters et al. 2021; Lamb 2021). Consequently, paralogons in many avian species, including chicken, are not split among chromosomes and, therefore, genes of a single paralogon are mostly kept on a single chromosome (Suppl. Fig. 1A). This characteristic enabled us to determine with high probability the chromosomal locations where missing avian ohnologs would belong.

For comparative analysis, the ohnolog genes of birds and several other clades of tetrapods were assembled into groups according to the mapping of their corresponding paralogons to chicken chromosomes. To avoid problems with poor annotation of some vertebrate genes, we searched only for ohnologs with orthologs discovered in the human genome, the most robustly annotated tetrapod organism. The analysis was initiated by comparing the ohnolog content of paralogons mapping to the chicken dot chromosomes (16, 29 – 32, 34 – 38) with the content of matching isoparalogons on other chromosomes (Fig. 1A, Suppl. Fig. 1B). Dot chromosomes were selected for the initial inquiry because of their high recombination rate as well as high repetitive content, making them a likely target for gene losses (Megens et al. 2009; McQueen, Siriaco, and Bird 1998). Ohnolog content in different clades of tetrapods was determined using the NCBI ortholog database as well as BLAST screens of genomes of selected species (Fig. 1B, Suppl. Tab. 1 and 2A-C). We observed only marginal loss of ohnologs in amphibians and two reptile lineages, lepidosaurs and turtles, and this loss was somewhat higher (2 – 3%) in paralogons that map in birds to dot chromosomes than in corresponding isoparalogons (0 – 1%). While losses in amphibians appeared taxon-specific, half of the genes lost in lepidosaurians were lost in all sauropsid taxons searched. In contrast to these marginal losses, birds have lost 29% of ohnologs from the paralogons mapping to avian dot chromosomes while only 0.4% of matching ohnologs have been lost from the other chromosomes. Almost the same number of ohnologs were missing from dot chromosomes of paleognaths and neognaths, and the missing sets of genes had substantial but not complete overlap. Interestingly, we noticed an increased number of missing genes in crocodiles, the reptile group most closely related to birds. In crocodiles, 13% of ohnologs mapping to the paralogons that are located in birds on dot chromosomes were not found. A majority of genes lost in crocodiles were lost in birds as well.

**Fig. 1.**
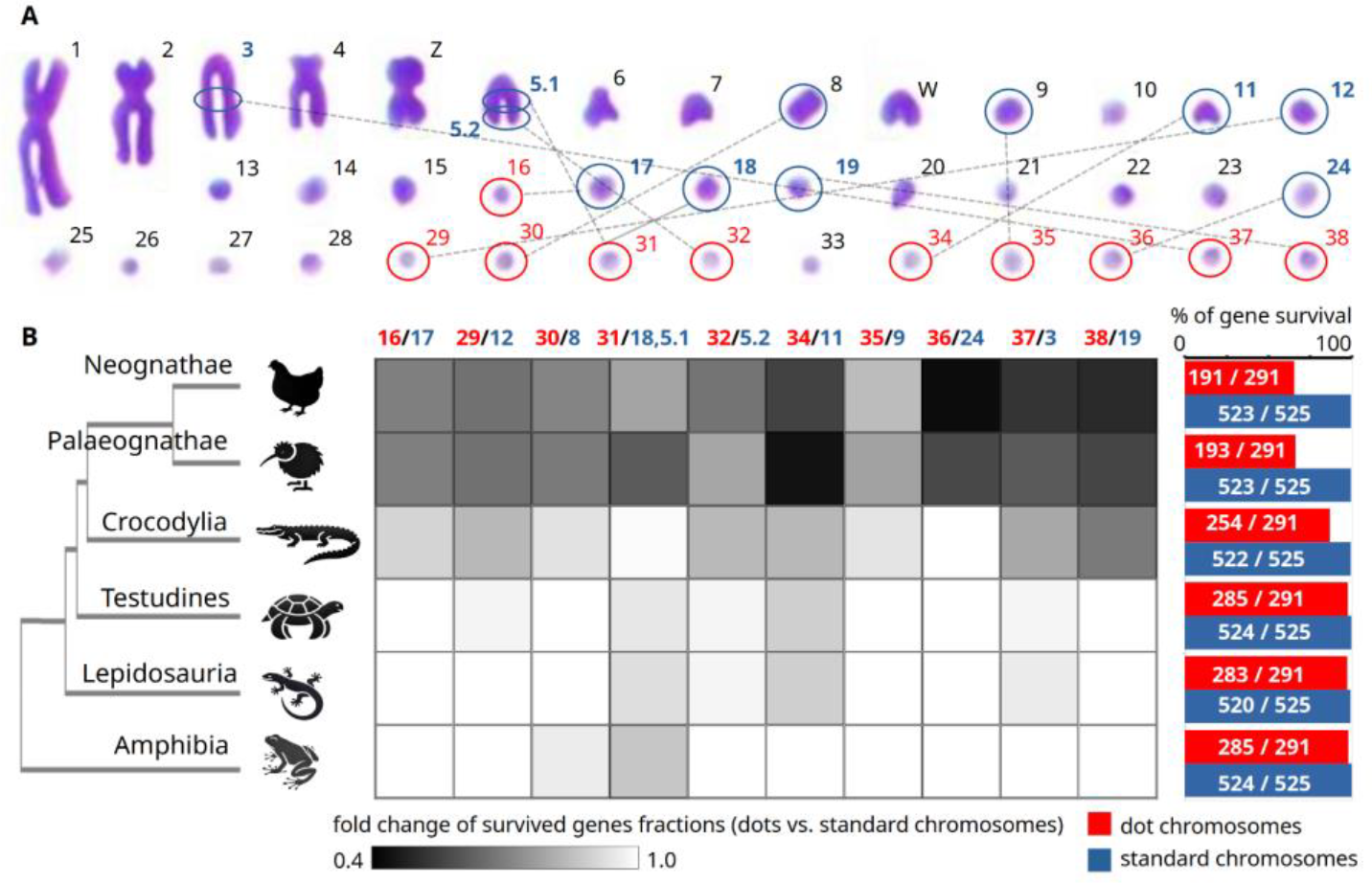
Evolutionary loss of genes that syntenically map to chicken dot chromosomes. Ohnologs from paralogous syntenic groups of genes (paralogons) that originated in the second whole genome duplication in the vertebrate ancestor were used to evaluate gene losses specific to chicken dot chromosomes. (A) Chicken karyotype of metaphase chromosomes adapted and modified from (Ji et al. 2016). Dot chromosomes are highlighted by red circles. Chromosomes or their parts carrying isoparalogons corresponding to the paralogons that map to chicken dot chromosomes are highlighted by blue circles and connected by dashed lines. (B) Number of ohnologs belonging to paralogons residing on chicken dot chromosomes and ohnologs in corresponding isoparalogons on other chromosomes were counted for each taxon. Counts of genes were then joined for each chromosome. Only ohnologs that have human orthologs were considered and the number of these genes in the human genome, therefore, represent the total number of genes counted in each paralogon. Counts of ohnologs in each vertebrate taxon shown represent the number of genes that survived during the evolution in that taxon. (B, left margin) The phylogenetic tree scheme indicates which row of results (heat map in the middle and bar plots in the right margin) represents results for each individual taxon. (B, right margin) Bar plots show percentages of surviving genes in all paralogons that map to dot chromosomes (red) and in their isoparalogon counterparts on other chromosomes (blue). (B, middle) Ratios of the surviving ohnolog fractions on each dot chromosome and their isoparalogon counterparts on other chromosomes are shown as a heat map.

To determine the gene losses on dot chromosomes in the principal avian model species, the chicken, we used the recently released genome assembly Ggswu of the chicken huxu breed (Huang, Xu, Bai, et al. 2023). This assembly is considered to contain a complete, telomere-to-telomere genomic sequence, except for chromosome W. Its quality is also supported by the detection of all previously reported “hidden” genes that were absent in the previous chicken genome assemblies. In addition to 85 genes that were missing in all bird species, BLAST searches in Ggswu failed to identify another 42 ohnologs from dot chromosome-mapping paralogons (Suppl. Tab. 2D). All of these additional genes were also missing in the ortholog database in other galliform species and two-thirds of them in anseriform species as well. In contrast, only two additional chicken genes in matching isoparalogons on other chromosomes were not identified by BLAST searches (Suppl. Tab. 2E). Galliform-specific gene losses suggest that the process of elimination of genes from dot chromosomes continued during the diversification of contemporary avian species.

Dot chromosomes are highly compartmentalized with distinct regions of euchromatin and heterochromatin (Huang, Xu, Bai, et al. 2023). Euchromatin is the region where most single-copy genes are located (meaning single-copy on a specific chromosome and ignoring paralogs on other chromosomes). In contrast, regions of heterochromatin neighboring the euchromatin are usually filled with a few classes of highly repetitive genes, such as olfactory receptors, zinc finger proteins, etc. While it is not possible to determine where lost genes would be located if they had survived, we mapped all ohnologs found on chicken dot chromosomes to the huxu genome, and all are located in euchromatin (Suppl. Fig. 2). That makes euchromatin the most probable region from which the genes were lost.

In a few cases the ohnologs expected to be found on dot chromosomes in synteny with other genes belonging to the same paralogon were found in non-syntenic positions on different chromosomes than expected and surrounded by genes from an unrelated paralogon. Positional relocations of the ohnolog IRF9 into six different non-syntenic positions, each in a different avian order, were described previously (Ungrová et al. 2025). Other positional relocations found during the course of this study include transfer of CARMIL3 from chr34 (paralogon 10A) to chr1 (5A6A) in Galliformes, PLEKHA4 from chr31 (4D6D) to chr15 (2D) in Neognathes, and ancient transfer of TPPP2 gene from chr34 (10A) to chrZ (7A8A) in ancestors of Archelosauria.

Dot chromosomes include the seven smallest chicken chromosomes. All ten dot chromosomes range in size, however, from 2.5 to 6.8 Mb and other non-dot microchromosomes fit within this range as well. To explore whether these other small chromosomes could also represent sites of preferential gene losses in birds, we compared the ohnolog content of paralogons mapping to all non-dot chromosomes smaller than 7 Mb with ohnologs in isoparalogons on other chromosomes for various avian species (Suppl. Fig. 1C). In all cases, the losses of avian genes were minor (Suppl. Fig. 3, Suppl. Tab. 2F). The highest losses were found on chromosomes 22, 25, and 33 (4-7%) and these losses were higher than in corresponding isoparalogons (0-3%). Chromosomes 25 and 33, while not classified as true dot chromosomes, have some similarities with dot chromosomes (Huang, Xu, Bai, et al. 2023). All other small microchromosomes tested (23, 25-28) lost less than 2% of ohnologs while matching isoparalogons did not lose any genes. These results eliminated avian chromosome size as a major factor determining the regions of observed gene losses.

Finally, we also tested the content of the remaining 20 paralogons that map to larger microchromosomes and to macrochromosomes (Suppl. Fig. 1). Only six ohnologs out of 800 (0.75%) belonging to these paralogons were not found in avian species in the ortholog database and were not detected by BLAST in chicken and ostrich genomes (Suppl. Tab.2G).

All these results suggest that up to 4% of genes from the 2766 ohnologs surveyed could be lost from all avian species. The presumptive losses are highly concentrated on dot chromosomes that lost on average 29% of ohnologs, with some of them losing almost 50%. Dot chromosomes in birds, as well as syntenic regions in other vertebrate species appear, therefore, prone to gene loss. Some losses might have already started in the ancestors of contemporary sauropsids, substantially increased in ancient archosaurs, and accelerated even more in the dinosaurian line leading to birds. The losses proceeded also to considerable extent in the descendants of the last common avian ancestor. Genes were most likely lost specifically from euchromatin regions of dot chromosomes where surviving syntenic ohnologs are located.

### Genes on dot chromosomes undergo massive local repeat expansion

Despite substantial gene loss, chicken dot chromosomes are known for being gene-rich (Huang, Xu, Bai, et al. 2023; J. Smith et al. 2000). We observed that genes present on dot chromosomes contain highly repetitive introns. Each gene intron usually contains one or several locally expanded motifs, resembling tandem repeat expansions; however, in these cases, repeated blocks are longer (tens to hundreds of bases) and less organized than in common microsatellite repeats. This can be clearly seen in sequence similarity dot plots, where one sequence is compared to itself using the BLASTn algorithm (Camacho et al. 2009) and homologous regions are plotted. Filled rectangular areas in the plots then represent regions of locally expanded repeats. Interestingly, when such repetitions were observed in ohnologs located on dot chromosomes, they were not present in their counterparts in matching isoparalogons that reside on other chromosomes. This suggests that repeat expansion occurred during or after the vertebrate WGD specifically on dot chromosomes. A representative example is gene PHF8 located on dot chromosome 29 and its ohnolog counterpart PHF2 located in a matching isoparalogon on standard chromosome 12 (Fig. 2A, and Fig. 2C upper schemes). Blocks of repeated sequences are isolated inside individual introns and we did not observe any case where they cross over to adjacent exons. Gene coding regions therefore remain intact.

**Fig. 2.**
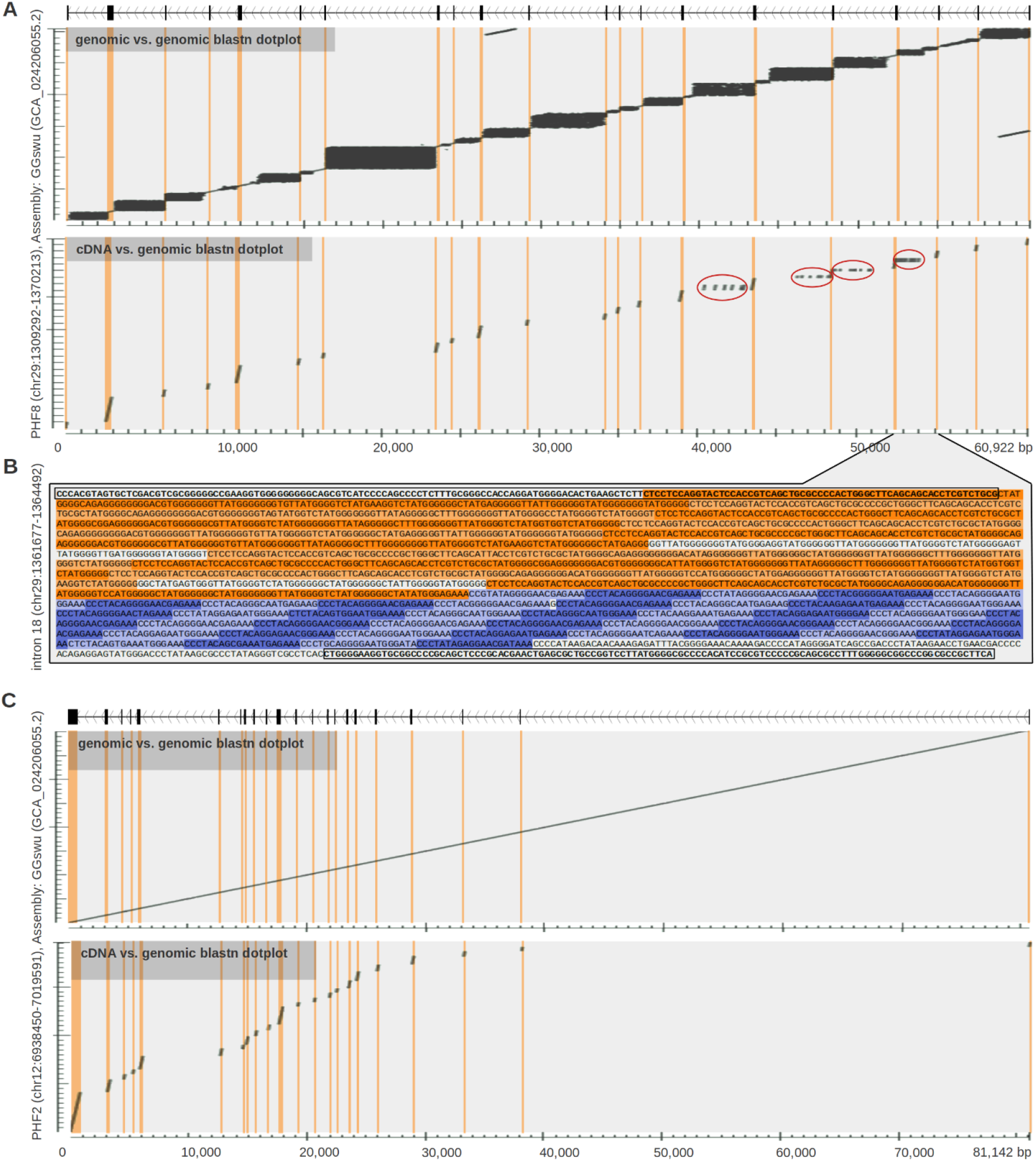
Example of sequence stuttering in avian gene introns. (A, C) Sequence dot plots show BLASTn alignments of genomic sequence vs. itself (upper schemes), and cDNA vs. genomic sequence (lower schemes). Dark regions in the plots represent BLASTn hits. Exon positions are manually curated and indicated in dot plots by orange vertical lines. (A) PHF8 gene located on chicken dot chromosome 29. Black rectangular regions in the upper dot plot indicate repetitive clusters usually mutually isolated by adjacent exons. Red circles in the lower dot plot indicate multiple copies of the sequence that originated in the closest adjacent exon. (B) DNA sequence of PHF8 intron 18. Adjacent exons are marked by black rectangle borders. Orange color marks one local repeat expansion involving a part of the left exon (exon stuttering). Blue color marks another local repeat expansion exclusively of intron origin (intron stuttering). Alternating lighter and darker orange and blue colors indicate individual repeated sequences. Repeats are defined as sequences with <20% edit distance from their consensus. (C) PHF2 gene located on chicken chromosome 12. This gene has originated from the same gene ancestor as PHF8 during the second WGD and shows no marks of sequence repetition.

In a subset of genes, we also observed a situation where part of the coding exon becomes the origin of the repeat cluster - e.g. part of the exon belongs to a repeated sequence block. Although this does not alter the exon region itself, it copies a lateral part of the exon, including the adjacent splice donor or acceptor site, through the intronic region and creates potential alternatively spliced transcripts. These repetitions were detected in PHF8 introns and are shown in a BLASTn dot plot of the genomic region versus a virtually spliced cDNA sequence but are not present in introns of the PHF2 gene (Fig. 2A, and Fig. 2C lower schemes).

Combination of two independent repeat expansion clusters in intron 18 of the PHF8 gene that originated from both exon and intronic sequences is demonstrated in the Fig. 2B. Manual inspection of the sequence showed that repeats are imperfect (<20% edit distance from consensus in this case) and structured in imperfect tandems. While it is important to note that some local variation can be caused by technical artifacts during sequencing and assembly processing, the general structure of tandem repeats in introns of PHF8 was independently verified by raw nanopore sequencing data. Observed expansions of originally unique sequences, as coding exons usually are, imply that repeat clusters have in general a local origin. Based on the repetitive “stuttered” pattern, we denoted this phenomenon as “sequence stuttering”. This term refers both to repeats exclusively of intron origin (intronic stuttering) as well as repeats involving also exon regions (exon stuttering). Other detailed examples of stuttering genes and their ohnolog counterparts from matching isoparalogonscounterparts from matching isoparalogons are shown in Suppl. Fig. 4.

We then aimed to systematically describe these repetitions in all chicken genes. In the majority of cases, sequence stuttering is recognised as a simple repeat region by an in-silico Tandem Repeats Finder screen (https://tandem.bu.edu/trf/home) (Benson 1999). We therefore analyzed the content of simple repeats for introns in individual chromosomes of the chicken genome assembly Ggswu (Fig. 3A). Indeed, dot chromosomes are clear outliers in intronic repeat content compared to the rest of the chicken chromosomes. This is consistent with previous observations where chicken dot chromosomes were shown to be highly repetitive (Huang, Xu, Bai, et al. 2023; Guizard et al. 2016).

**Fig. 3.**
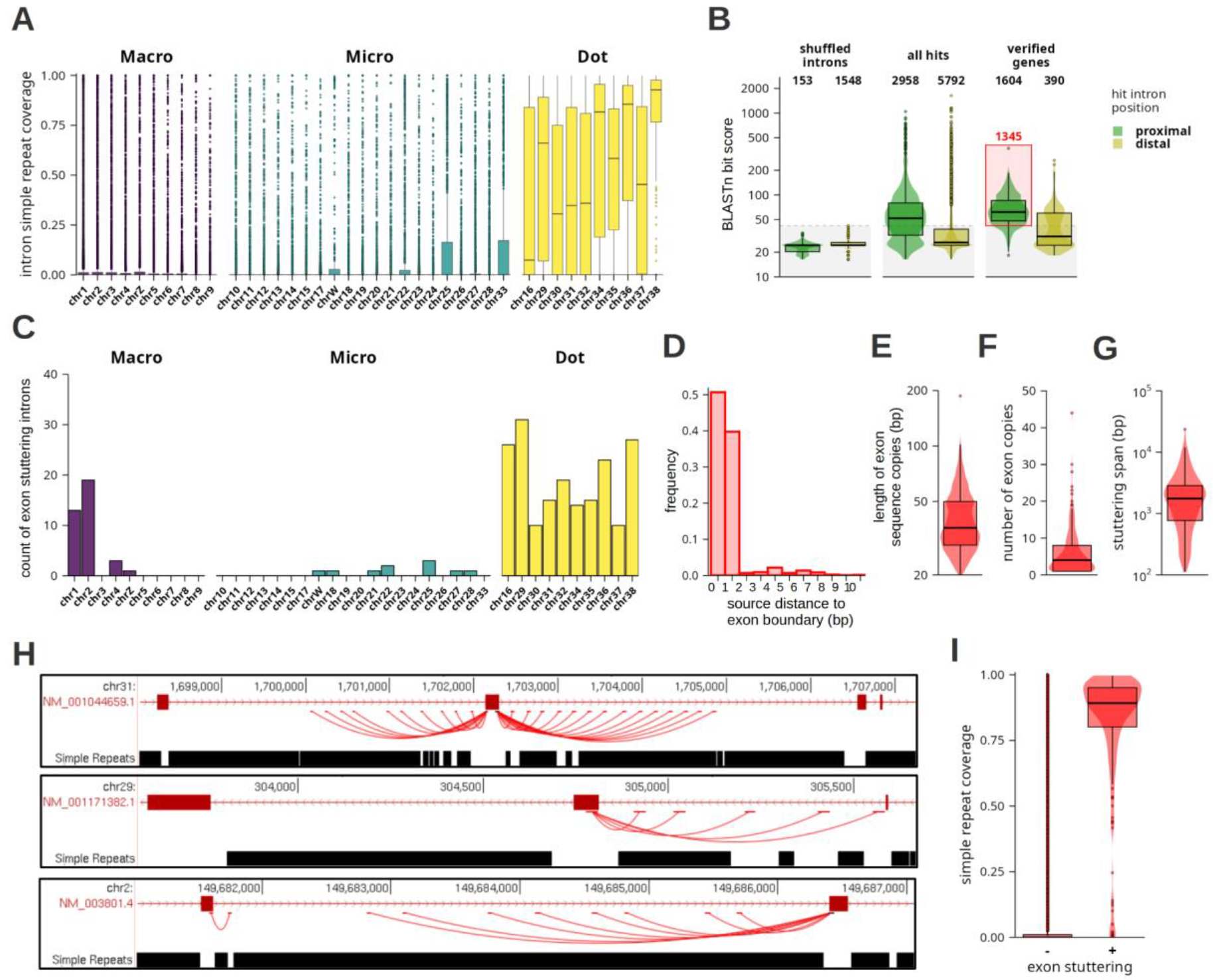
Repeat expansion and exon stuttering in the chicken genome. Data are presented as a box plot - panel A, combination of box and violin plots - panels B, E-G, I, bar graphs - C-D, or a BED type graph - H. In box/violin plots the horizontal line inside the box indicates median, lower and upper ends of the box indicate 1^st^ and 3^rd^ quartile, whiskers represent a data range spanning percentiles 0.05 to 0.95, and dots represent individual introns outside the range. (A) Distribution of intron coverage by simple repeats. Introns are grouped by chromosomes. Colors indicate chicken Macro-, Micro-, and Dot-chromosomes. (B) Results of exon stuttering screen in the chicken genome. Bit score distributions of BLASTn hits of exon sequences in all chicken introns are plotted for proximal and distal introns separately. Distributions are shown for: shuffled introns - hits in control shuffled intron sequences, all hits - unfiltered hits in real intron sequences, verified genes - hits present only in a set of genes whose genome annotations were manually verified. Red rectangle marks hits that passed all filtering criterias (proximal hits, bit score above shuffled control, present in verified genes) and are analyzed in the following panels C-I of this figure. (C) Number of introns with exon stuttering (introns containing one or more partial copies of adjacent exon sequences) calculated for each individual chicken chromosome. (D) Distance distribution of copied exon source sequences from the exon border. (E) Distribution of lengths of individual exon sequence copies. (F) Distribution of number of exon sequence copies for individual expansions. (G) Stuttering span distribution of individual exon expansion regions. Stuttering span is defined as a length of genomic region spanning all copies of a particular exon sequence. (H) Representative examples of exon stuttering for 3 different chicken genomic loci. Upper track shows XenoRefSeq gene annotation (original UCSC track), middle track shows exon sequence copies where each line connects source exon sequence with individual copies in adjacent introns (confident BLASTn hits from our screen), lower track shows content of simple repeats (original UCSC track). (I) Distribution of the simple repeat coverage for all chicken introns compared to exon stuttering introns.

To identify stuttering genes more specifically, we performed a homology-based screen evaluating multiplication of exonic sequences (exon stuttering) in chicken gene introns (Fig. 3B, Suppl. file 1). In this screen, sequence hits of exonic sequence occurring in neighbouring introns were referred to as proximal, whereas hits occurring in other introns were referred to as distal. Noise represented by false positive hits was estimated by a shuffled control where the screen was performed on randomly shuffled intron sequences (Fig. 3B, shuffled introns). As expected from the random control, the vast majority of the hits were distal. The median BLASTn bit score of random control hits was 24.3 and the maximal value was 42.1. We, therefore, used the bits core 42.1 as a confidence threshold for the screen results. Because the sequence stuttering occurs locally in clusters, we expected that true screen hits would be located in proximal introns. Indeed, a screen of chicken data showed 2,958 hits in proximal introns with a median bit score of 52 (Fig. 3B, all hits). The remaining 5,792 hits were localised in distal introns, however most of them aligned to the source exonic sequence only poorly (median bit score 26.3, 78% of hits below bit score 42.1) and represented mainly false positive hits. To further increase the specificity of the screen, we manually reviewed all 192 genes with significant proximal hits and excluded 77 of them as false positives, mainly because of incorrect gene annotation. Finally, we obtained 115 genes containing 1,345 confident hits comprising 236 introns with putative repeat expansion of exonic sequences (Fig. 3B, verified genes).

190 out of 236 identified introns were located on dot chromosomes, 10 on other microchromosomes, and 36 on macrochromosomes (Fig. 3C). Considering that dot chromosomes contain less than 4% of all chicken introns, this result suggests an extremely strong preference for exon stuttering introns to be located on dots. This screen in principle identified only stuttering introns where a substantial part of the exon sequence is copied. Expansions of purely intronic sequences were thus not recognized and the real number of stuttering introns is expected to be much larger.

Source sequences of individual exon expansions were in more than 90% of cases located a maximum of 1 base from the exon border (Fig. 3D). This suggests that the majority of copied regions span an exon-intron border and contain part of the intron sequence. The exon parts of the expanded sequences (exon hits) had lengths from 20 to 187 bp with median 36 bp (Fig. 3E). Although the full lengths of expanded sequences (including intronic regions) cannot be determined without manual curation, these results seem to be consistent with our previous observations that lengths of repeated blocks range from tens to hundreds of bases. We detected from 1 to 44 copies (median 4) in individual expansions (Fig. 3F) and the lengths of whole expansion areas, including the source sequence, were 112-23,555 bp (median 1,751; Fig. 3G). Cases where we identified only a single copy of the exon sequence may represent an early stage of sequence expansion or they may eventually be a result of other genomic processes. Manual inspection of exon stuttering hits showed that they generally exhibited a prototypic pattern of sequence stuttering as described earlier in the manuscript (Fig. 3H). Consistent with this, exon stuttering introns were also heavily covered by predicted simple repeats (Fig. 3I).

In summary, we identified an unusual phenomenon—sequence stuttering—that is at least partially responsible for the high repetitiveness of sequences on dot chromosomes. Observed repeat expansions are strictly local, meaning that they are isolated between evolutionarily conserved coding sequences - gene exons. Analysis of gene introns was therefore shown to be a very valuable method for describing this process. An automated screen of sequence stuttering events is very difficult due to the high variability in the noncoding regions and the complex pattern of expanded regions. We therefore developed a screening strategy that is capable of identifying specific cases of sequence stuttering where part of the coding exon sequence is expanded. The screen confirmed that this process is tightly connected to dot chromosomes. Considering that this screen identified exon stuttering in 190 out of 5,572 (3.4%) annotated dot chromosome introns, it indicates that at least this percentage of all stuttering introns is expected to contain detectable exon sequence copies. The method, therefore, seems to be valuable in identifying sequence stuttering phenomena and can potentially be used for large-scale screens in future work.

### Dynamic nature of local repeat expansion in chicken population

Although dot chromosome regions containing stuttering genes are largely missing in avian genome assemblies, in some instances we observed local repeat expansions also in other birds besides chicken. Comparing two orthologous introns in closely related species, we noticed that they often have different lengths and contain repeat expansion clusters of differing structure and sequence. This suggests that repeats expanded in individual species independently.

To understand whether intron variability extends also to the intraspecies level, we analyzed public nanopore sequencing datasets of different chicken breeds and compared the sequence and structure of selected stuttering gene introns. First, we retrieved the reads containing complete sequences of individual introns. Because identification of reads based on intron reference sequences failed due to low sequence complexity, high variability and low sequencing quality of intronic regions, we identified intron-containing reads using adjacent (evolutionarily conserved) exons. Interestingly, we observed that stuttering introns are highly variable in their lengths both among and within individual samples. As a case example, we show intron 18 of the aforementioned stuttering gene PHF8 (Fig. 2B, 4A). This intron contains two distinct repeat clusters, visualized as rectangular regions in a BLASTn dotplot: one originating partially from exon 19 (Fig. 4A, left exon) and another located completely inside the intron. Nanopore read data showed that this intron varies in length from 1.6 to 2.9 kb and, in some individuals, there is a prominent pattern of two distinct-length alleles, indicating heterozygosity (Fig. 4B).

**Fig. 4.**
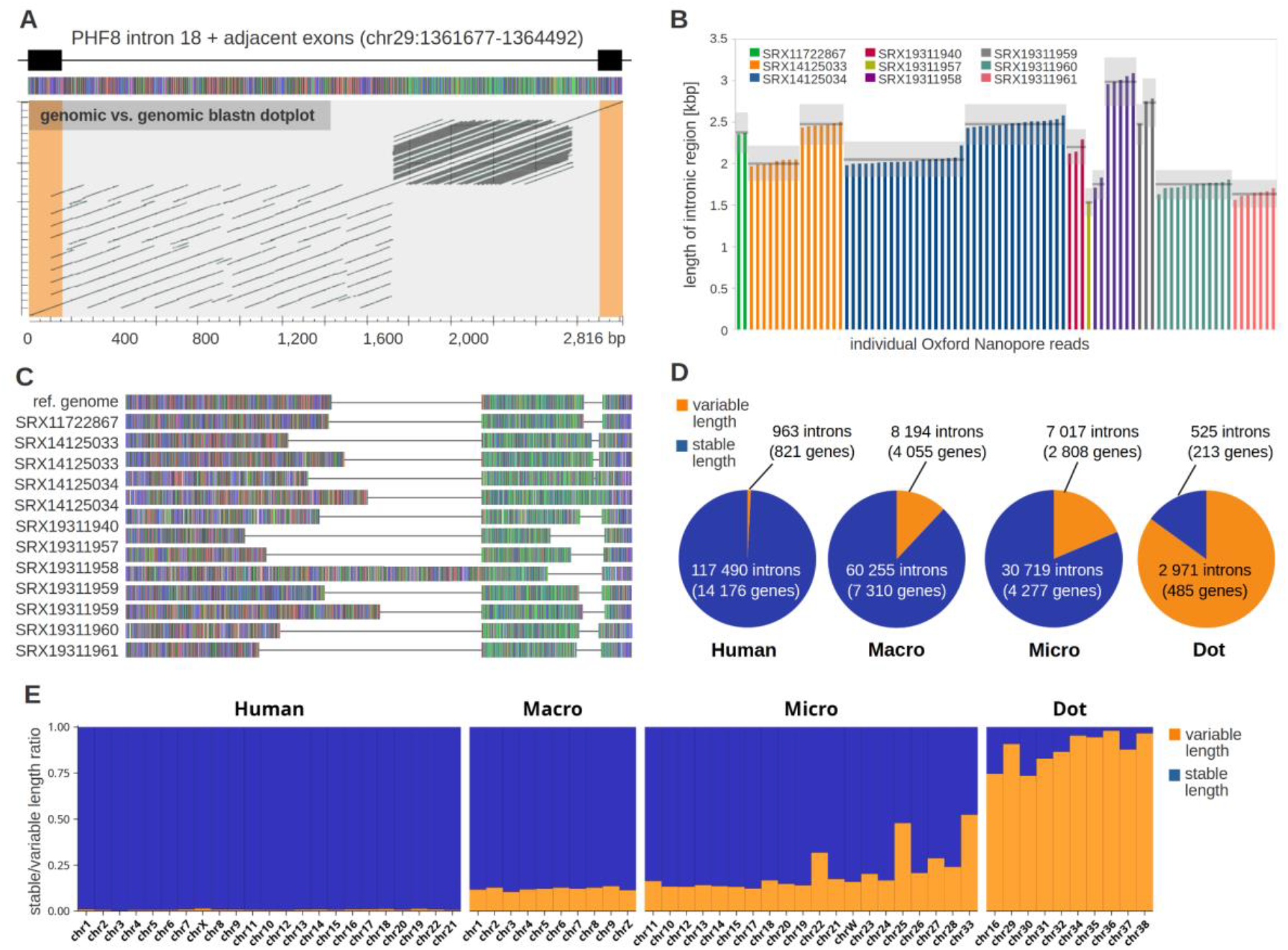
Intron length variation is concentrated on chicken dot chromosomes. (A) PHF8 intron 18 as an example of chicken “stuttering intron”. The GGswu reference sequence with positions of the exons and the intron is indicated at the top and the colored scheme of sequence is shown below. Each nucleotide is represented by a different color (A - green, C - blue, G - grey, T - red). Sequence dot plot at the bottom shows BLASTn self-alignments of genomic sequences, where dark diagonals represent BLASTn hits. Exon positions were manually curated and are indicated inside the dot plot with orange rectangles. (B) Chicken PHF8 intron 18 length was determined in public Nanopore sequencing data. Bars in the plot represent intron lengths estimated from discrete nanopore reads. Colors of the bars indicate individual datasets (individual chickens). Datasets are also identified by sequence accession numbers from NCBI SRA stores. Intron length clusters representing alleles in individual samples are indicated by horizontal dark grey lines (light grey areas show +/-10% intervals from average length of each allele). (C) Schematic representation of allele consensus sequences of PHF8 intron 18 identified in public Nanopore sequencing data. Colors represent individual nucleotides as in panel A. Sequences are aligned based on the positions of two repeat clusters shown in panel A dotplot. Horizontal lines in each sequence scheme represent gaps in the alignment. (D) Identification of length variable introns in human and chicken populations. Intron length alleles were identified in various human and chicken Nanopore sequencing data. Pie plots show the number of introns in human genome (Human), chicken macrochromosomes (Macro), chicken microchromosomes (Micro), and chicken dot chromosomes (Dot) for the following categories: stable (no length variation in the samples - rarefied allele richness = 1), variable (more than one length allele in the samples - rarefied allele richness > 1). (E) Fractions of length variable and stable introns for individual human and chicken chromosomes.

Then, we clustered intron sequences from nanopore reads into individual alleles and created their consensus sequences (Suppl. file 2). Problematic sequences of these introns and the consequent high error rate of sequencing data precluded reconstructing exact consensus sequences in sufficient quality for detailed short nucleotide variation analysis. Therefore, we instead compared the overall length of each allele and its repeat structure based on the BLASTn dotplot patterns (Suppl. Fig. 5). Indeed, the variable length of alleles was shown to be caused by expansion or reduction of individual repeat clusters. This point can be demonstrated when alleles from various chicken samples are aligned based on the positions of two repeat clusters occurring in the intron (Fig. 4C).

To assess intron length variation across the entire chicken genome we developed an automated screen, following the approach described above for PHF8 intron 18. We analyzed public nanopore datasets of 11 individual chickens and 3 humans as a control (Suppl. files 3 and 4). Due to occasional inaccuracies of gene annotation or extreme sequence properties, some genomic regions can yield false results. To eliminate such cases, we built a set of filtering criteria described in the Materials and Methods. After filtering, we obtained informative results for 70% of all annotated chicken introns and for 49% of human introns. Lengths of individual human and chicken introns were clustered into length alleles. Distinct alleles were defined as sets of intron sequences whose mean length differed from other alleles by more than 10%. This criterion was used to exclude false-positive alleles produced by sequencing indel errors. The numbers of distinct length alleles for individual introns were estimated and rarefied to 3 diploid samples (n=6). This metric was further used as an estimation of intron length variability in the population and was referred to as rarefied allele richness (Ar). Value 1 means that 3 random individuals from the chicken population are expected to all contain a given intron of the same length, whereas value 6 means that they are expected to contain all 6 intron copies with different lengths. Introns with Ar = 1 were considered as stable, whereas introns with Ar > 1 were considered as length-variable.

Screening of human data revealed that less than 1% of introns are variable (Fig. 4D) and 88% of variable genes contain only a single variable intron (human variable genes contain an average of 1.2 variable introns). This reflects an expected situation where variability in intron length is caused by occasional genetic events, such as mobile element insertion or tandem repeat expansion. Because human and chicken populations have very different sizes and structures, comparison of intron length variation between them is not very informative. Human results, therefore, serve as a proof of principle of the method and for basic comparison.

Chicken samples yielded 12% and 26% of variable introns on macrochromosomes and microchromosomes excluding dots, respectively (Fig. 4D). This contrasts sharply with the 85% of variable introns observed on dot chromosomes (Fig. 4D). Additionally, 87% of the variable intron-containing genes on dot chromosomes have more than a single variable intron (an average of 6.1 variable introns). Examination of individual chromosomes confirmed that this feature is common for all chicken dot chromosomes (Fig. 4E). Slightly elevated levels of variable introns were observed also in chicken chromosomes 22, 25, and 33. Interestingly, as noted earlier, these chromosomes were also shown to have the highest rate of gene losses, besides all dot chromosomes. These analyses revealed exceptional evolutionary dynamics in the chicken population concentrated on dot chromosomes.

In the chicken samples, we identified 18,183 length variable introns, where 2,971 of them reside on dot chromosomes (Fig. 5A). Although this represents only a fraction of all variable introns, it covers the vast majority of introns located on dots (only 522 dot chromosome introns were identified as stable). The source of length variation for at least a subset of introns seems to be sequence stuttering, as we described earlier. Consistent with this, 95% of introns (173 out of 183) where we identified exon sequence stuttering (screen described in Fig. 3) are also length-variable. Moreover, the estimated level of population variability of exon-stuttering introns is exceptionally high. The median of Ar for exon-stuttering introns is 3.9, compared to the nearly two-fold lower median of all variable introns, 1.9 (Fig. 5 B). Introns located on dot chromosomes in general also exhibit substantially higher variability (median Ar 2.7) compared to the variable introns’ median. Besides the measure of population variability, we also inspected absolute differences in lengths of individual intron alleles in a single representative chicken individual (Fig. 5C). The majority of variable introns differ in length only slightly (median difference 19 bp). In most cases, this is probably caused by common genetic variability, including short indel variants. Substantially larger differences were observed in introns located on dot chromosomes (median difference 68 bp). This applies especially to introns where exon stuttering was identified (median difference 549 bp). In these cases, we observed that variation in intron lengths is generally caused by the dynamics in individual copies of stuttering regions.

**Fig. 5.**
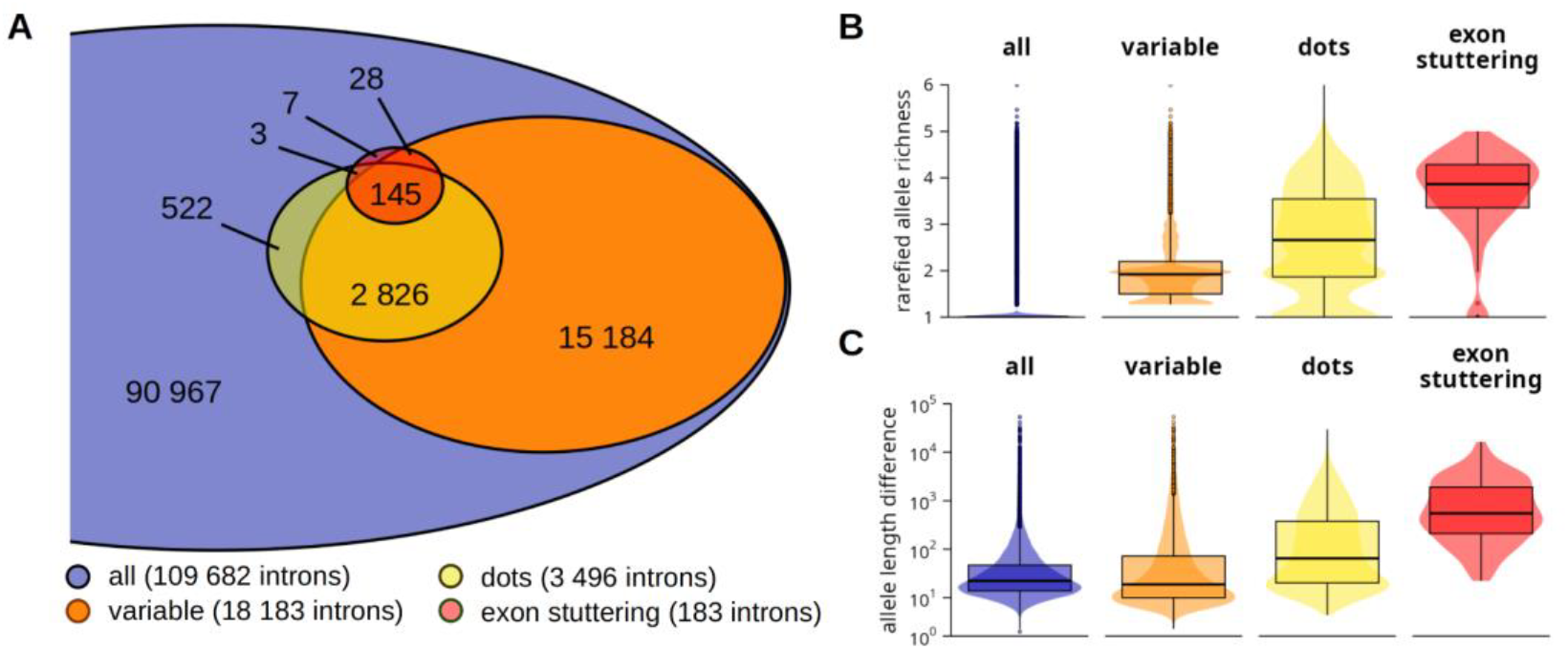
Extensive length variation is characteristic for chicken stuttering genes on dot chromosomes. (A) Venn diagram of chicken introns that passed the length variation screen. Numbers indicate counts of introns in each intersection. variable - introns with rarefied allele richness > 1, dots - introns located on dot chromosomes, exon stuttering - introns identified by the exon stuttering screen. (B) Rarefied length allele richness Ar of chicken introns that passed the length variation screen grouped by categories defined in section A. (C) Absolute length difference between alleles of chicken introns that passed the length variation screen. Differences in base pairs were calculated for selected chicken samples (SRX14125033) and grouped by categories defined in section A. (B, C) Whiskers of the box plots represent a data range spanning percentiles 0.05 to 0.95, dots represent individual introns outside the range.

Taking together, we revealed that exon stuttering introns identified by our screen and found prominently on dot chromosomes manifest exceptional population variability. Also, random inspection of other introns located on dot chromosomes confirmed that introns showing sequence stuttering properties are highly variable in general (examples are shown for several stuttering genes and their isoparalogon ohnologs in Suppl. table 3).

## DISCUSSION

In this study, we describe two genetic processes strongly associated with avian dot chromosomes: gene loss through avian evolution and a novel form of genetic instability characterized by local expansion of genomic sequences.

Gene loss is a common driver of genetic variation across all organisms, occurring either randomly or with adaptive significance (Albalat and Cañestro 2016). The absence of a substantial number of evolutionarily conserved genes in the genome of avian species is a long-time unsolved cardinal question of avian genetics. Elucidation of this problem is complicated not only by avian hidden genes (Hron et al. 2015) but also by the considerable number of gene paralogs (ohnologs) present in the avian genome, which originated during whole genome duplication (WGD) in vertebrate ancestors (Singh and Isambert 2020; Huang, Xu, Bai, et al. 2023). We have observed that various genes, for example, IRF3 and IRF9, which were previously claimed to be found in chicken, were actually confused with their paralogs (Yin et al. 2019; Ungrová et al. 2025). Most of these errors were due to gene identification based solely on sequence similarity and not on gene synteny. Therefore, using the knowledge of the conserved synteny of ohnologs throughout all jawed vertebrate species (Simakov et al. 2020; Marlétaz et al. 2024; Lamb 2021), we explored the presence of a set of ohnologs in birds and compared it with presence of their orthologs in other tetrapod species. We have found that about 4% of ohnologs we searched for in birds were lost in all avian species, and 75% of these gene losses were concentrated on dot microchromosomes, which carry only about 5-10% of avian single-copy genes, i.e. genes not generated by segmental duplication. One hundred and twelve ohnologs missing in avian genomes represent likely only a fraction of genes lost. Preliminary rough estimates obtained by searching for all single-copy genes (not only ohnologs) that, based on their synteny in multiple vertebrate species, belong to paralogon 10A suggest by extrapolation that several hundreds of genes could be lost in all birds (data not shown).

Reliability of the determination of gene loss in birds depends on several factors. First, it is the number and completeness of avian genome assemblies available. Currently, almost 1,600 avian species out of more than 11,000 recognized have their genome sequenced and deposited in the NCBI database (https://www.ncbi.nlm.nih.gov/datasets/genome/). The quality of genomes has also improved. New assemblies generated by the Vertebrate Genomes Project now contain up to 11% of additional, often GC-rich sequences that were completely missing in previous assemblies (Kim et al. 2022). The second factor is the reliability of the NCBI eukaryotic genome annotation pipeline which generates the NCBI ortholog database (https://www.ncbi.nlm.nih.gov/kis/info/how-are-orthologs-calculated/) that we used extensively. Finally, all losses were verified by BLAST searches in the chicken Ggswu genome (Huang, Xu, Bai, et al. 2023). This is a telomere-to-telomere assembly and likely the most complete avian genome available. We detected only three known chicken genes missing from this assembly in the terminal region of chromosome 35. The correctness of gene loss was also in some cases corroborated by multiple nanopore datasets.

Seminal work on the loss of syntenic gene clusters in birds (Lovell et al. 2014) advanced the discussion about the losses of avian genes. The list of 274 genes presented as missing is 30% composed of ohnologs, mostly from dot chromosomes. We found approximately 50% of those genes, while 50% remain in the category of missing genes. Interestingly, not only ohnologs but a total of approximately 90% of all genes from the 274-gene list map to syntenic clusters on avian dot chromosomes. This partially explains the loss of syntenic blocks presented by the authors. We currently cannot exclude that, in addition to macrosynteny, there is also a loss of microsyntenic blocks, as this question will require further studies.

In a small fraction of cases, the ohnologs with an expected syntenic location on dot chromosomes (CARMIL3, IRF9, PLEKHA4, TPPP2) were not lost but relocated to noncanonical non-syntenic positions. Positional relocation of genes is a phenomenon described previously, see for example (Bhutkar et al. 2007). It would be interesting to determine if the frequency of relocation from dot chromosomes is higher than relocations from other regions of the avian genome and if it might represent an “emergency” measure to save important genes from being lost.

Loss of genes located on dot chromosomes was apparently a lengthy process. Because of the minimal gene loss in turtles and a very substantial loss in crocodylians (⅓ of genes lost in birds), the process must have started in the late Permian and did not end until some time after the split of galliform and anseriform birds in the late Cretaceous (Kumar et al. 2022). It took at least 200 million years in total. We don’t have, however, any definitive proof that the gene losses stopped, so it is possible that they are still continuing. While in agreement with a previous study (Lovell et al. 2014) the largest number of genes was likely lost before the last common ancestor of contemporary birds evolved, the entire process took an extended time and was not exclusively connected with the evolution of genetic traits of the class Aves.

Mechanisms of gene losses on dot chromosomes are not entirely clear. Because surviving ohnologs are found exclusively in euchromatin regions of dot chromosomes, it is most likely that those are also the regions from which genes were lost. Therefore, the driving force for loss appears to be chromosomal location, not the adaptive nature of loss. Based on the Ggswu chicken genome sequence, euchromatin regions occupy about half of each dot chromosome, are devoid of interspersed repeats, but are filled with GC-rich and simple tandem repeat-rich sequences with characteristics of sequence stuttering that are discussed in the following paragraphs. The absence of interspersed repeats, high gene density, and intraspecies variability of stuttering sequences suggest that this is a highly dynamic environment where only essential gene sequences could survive. This environment could also erase all traces of lost genes, as we did not encounter any pseudogenes left after lost genes. It is possible to imagine a step-by-step scenario where GC biased gene conversion acting on dot chromosomes leads to an increase in GC content of gene coding sequences and consequently to increased variability of encoded proteins (Botero-Castro and Wolf 2025; Huttener et al. 2021). This, in turn, may loosen the interaction of these proteins with their interaction partners, and genes with products not firmly anchored in an interaction network may then be more easily lost under appropriate circumstances. The function of lost ohnologs could be at least partially replaced by their paralogs (Drobek 2022).

The loss of genes from dot chromosomes invites the question: why do birds keep dot chromosomes instead of fusing them into larger units as is often the case in other sauropsids? Would the birds and in general the entire dinosaur evolutionary line that culminated by the class Aves benefited, for example, from the accelerated evolution of protein-coding genes on dot chromosomes? The presence of four hidden GC-rich muscle-specific genes ALDOA, ENO3, PYGM, SLC2A4 (GLUT4) on dot chromosomes (chicken chromosomes 38, 35, 37, 35, respectively) could indicate such a hypothetical benefit. As proposed by (Huttener et al. 2021), specific chromosomal locations of muscle-specific genes in GC-rich chromosomal regions that contrast with positions of their paralogs (ohnologs) with muscle-unrelated functions outside of the GC-rich genomic regions might suggest orchestrated accelerated protein evolution focused on special energy needs of avian musculature.

What we describe in this work as intron sequence stuttering is, to the best of our knowledge, a novel genetic phenomenon, distinct from known types of intron length variation (Fingerhut and Yamashita 2022; Bradnam and Korf 2008). Its three main features, namely intron repetitiveness, extensive intraspecies length variation, and occasional involvement of exon sequence fragments, are schematically depicted in Fig. 6. Currently, we describe the stuttering phenomenon in birds, and we did not find it in other vertebrate genomes. A notable exception is a group of genes identified in rodents (gerbils); these are extremely GC-rich (Hargreaves et al. 2017) and, according to our analysis, the pattern of sequence stuttering is also present in their introns (data not shown). This strengthens the potential association of sequence stuttering with the GC-rich nature of coding sequences. Further, although our analysis focuses mainly on gene introns, intergenic sequences on dot chromosomes are also affected by extensive variation of short repeats.

**Fig. 6.**
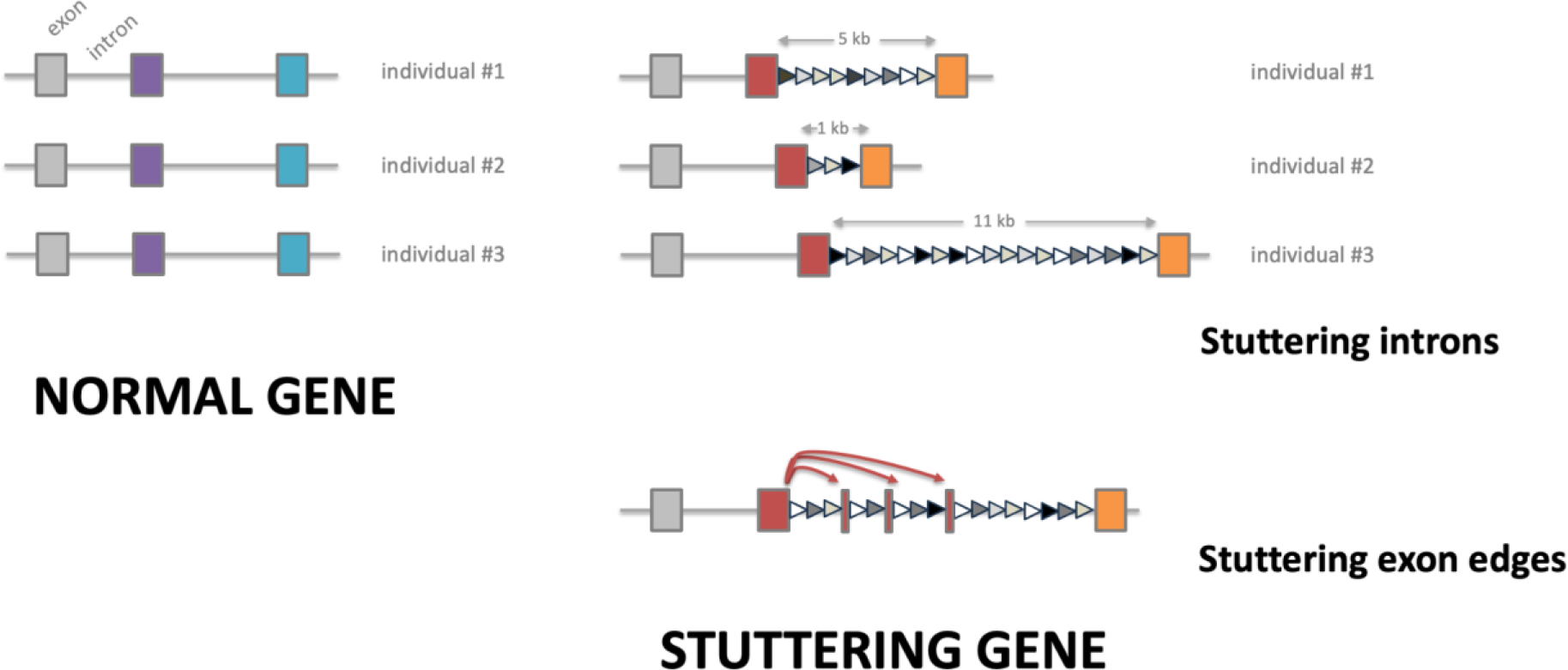
Sequence stuttering of avian genes. Schematic representation of three-exon hypothetical genes. Normal genes have uniform intron lengths in each individual chicken. Stuttering occurs in the second exon, characterized by imperfect short direct repeats (triangles of varying gray shades) forming introns with highly diverse lengths. Exon stuttering is represented as edge fragments of red exon interspersed multiple times into the neighboring intron.

At present, we can only speculate about the molecular mechanisms responsible for stuttering generation. One possibility is the involvement of mitotic and DNA repair processes, which might be reinforced in genomic loci under replication stress caused by high GC content or DNA secondary structures (Kruisselbrink et al. 2008; Kamp et al. 2021; Lemmens, van Schendel, and Tijsterman 2015). Indeed, multiple chicken hidden GC-rich genes were predicted to contain atypical DNA structures, mainly DNA quadruplexes (G4) (Beauclair et al. 2019; Hron et al. 2015). G4-enriched regions of the human genome have also been shown recently to be hotspots of genetic instability (Williams et al. 2020). An alternative mechanism might involve meiotic events, where inaccurate strand invasion during chromosome alignment in repetitive stuttering regions would be resolved as non-crossover gene conversion events. This would have the capacity to gradually increase or decrease the length of repetitive introns across generations, in a similar way to meiotic expansion/contraction mediated by tandem repeats (Jurka 2004; Sasaki, Lange, and Keeney 2010).

The results that show both phenomena, gene loss and gene sequence stuttering, in birds predominantly occurring on dot chromosomes, raise questions about the evolutionary origins of this special chromosome category. We believe that their origin could be traced to the second WGD. The second round of WGD in the evolution of vertebrates is assumed to be allopolyploidy, the rise of a polyploid species through interspecies hybridization (Simakov et al. 2020). The nature of this event was recognized based on biased fractionation, a phenomenon where two genomes contributed to a new species by parent lineages behave asymmetrically (Session 2024). One of the subgenomes (genomes contributed by diploid parents of a polyploid progeny) becomes dominant, leading to the preferential gene loss from the other, recessive subgenome. Reconstruction of the ancestral vertebrate karyotype suggested that genes of the recessive subgenome are located preferentially on avian microchromosomes rather than macrochromosomes (Huang, Xu, Cai, et al. 2023). A detailed comparison of the number of ohnologs identified in four vertebrate non-avian species (Lamb 2021) and assigned to paralogons mapping to dot chromosomes and to their isoparalogons (Supplementary Fig. 6) confirmed that all dot chromosomes appear to be descendants of chromosomes of the recessive subgenome. Dot chromosomes lost a disproportionate number of genes between the second WGD and the time when the last common ancestor of contemporary bony vertebrates (Osteichthyes) evolved, became the smallest chromosomes, and were likely a preferential target of frequent meiotic recombination and gBGC. While in the majority of vertebrates many of these smallest chromosomes fused together or to larger chromosomes, in the evolutionary line leading to birds they did not undergo any fusion (except chicken chr31, which is apparently the result of an ancient fusion of two small microchromosomes after the second WGD) (Waters et al. 2021; Huang, Xu, Bai, et al. 2023). We hypothesize that the accumulated evolutionary time when these chromosomes remained unfused and were subjected to frequent meiotic recombination and gBGC led to an increase in GC content, accelerated evolution of gene coding sequences, accumulation of repetitive content, and finally to increased gene losses. All these accumulated sequence changes then culminated in Archosauria and especially in birds, where a massive second wave of gene losses occurred.

In summary, our study reveals avian dot chromosomes as primary sites of extensive gene loss, with 29% of ohnologs absent, and a novel genetic instability, termed sequence stuttering, characterized by intronic repeat expansions and significant length polymorphism. These processes, likely driven by elevated GC content and recombination rates, provide important insights for future investigations into vertebrate genomic evolution.

## SUPPLEMENTARY DATA

**Supplementary figures 1-6**

**Supplementary tables 1-3**

**Supplementary file 1**: UCSC Genome Browser interact file containing BLASTn hits of exon stuttering screen. Score (5th column): bit score of BLASTn hit, exp (7th column): category of a hit based on our filtering criteria, sourceName (12th column): exon id, targetName (17th column): intron id containing exon ids of adjacent exons. **Supplementary file 2**: Sequences of manually constructed alleles of chicken PHF8 gene, intron 18, in fasta format. **Supplementary file 3**: Table of chicken introns based on the XenoRefSeq annotation of GGswu genome assembly. Columns contain various data described in this study.

**Supplementary file 4**: Table of human introns based on the Ensembl annotation of GRCH38 genome assembly. Columns contain various data described in this study.

## MATERIALS AND METHODS

### Reference genomes and gene annotations

For analysis of the human genome, reference assembly GRCh38.p14 (GCA_000001405.29) and ENSEMBL gene annotation version 112 were used. For analysis of the chicken genome, assembly GGswu (GCA_024206055.2) was used. As no official gene annotation for this assembly is available, RefSeq mRNAs Track (XenoRef, GCA_024206055.2_GGswu.xenoRefGene.gtf.gz, version 2024-12-03) present in the UCSC Genome Browser (UCSC GB) (Perez et al. 2025) hub directory (Clawson et al. 2023) was used in this work. The description of this annotation track, as is mentioned in the UCSC GB, is the following: “The mRNAs were aligned against the Gallus gallus/GCA_024206055.2_GGswu genome using translated blat. When a single mRNA aligned in multiple places, the alignment having the highest base identity was found. Only those alignments having a base identity level within 1% of the best and at least 25% base identity with the genomic sequence were kept. Specifically, the translated blat command is: blat -noHead -q=rnax -t=dnax -mask=lower target.fa query.fa target.query.psl where target.fa is one of the chromosome sequence of the genome assembly, and the query.fa is the mRNAs from RefSeq. The resulting PSL outputs are filtered: pslCDnaFilter -minId=0.35 - minCover=0.25 -globalNearBest=0.0100 -minQSize=20 -ignoreIntrons -repsAsMatch - ignoreNs -bestOverlap all.results.psl GCA_024206055.2_GGswu.xenoRefGene.psl. The filtered GCA_024206055.2_GGswu.xenoRefGene.psl is converted to genePred data to display for this track.”

This chicken gene annotation is not accurate in the exact exon borders and fails to annotate some genes entirely. However, despite these disadvantages it is still relatively complete. For detailed analyses of genes PHF8, PHF2, AKT1, AKT2, LFNA, and LFNB, the UCSC annotation was corrected by inspection of various publicly available RNA seq data.

### Analysis of gene ohnologs

Lists of ohnologs and the affiliation of each ohnolog to a specific paralogon from 68 paralogons recognized were published previously (Lamb 2021). Paralogons were assigned to chicken chromosomes based on the data published by (Huang, Xu, Bai, et al. 2023).

Presence of ohnolog genes in genomes of all species in the NCBI databases belonging to selected vertebrate taxons was determined by querying NCBI databases by gene symbol and taxon using datasets command-line tool (https://www.ncbi.nlm.nih.gov/datasets/docs/v2/reference-docs/command-line/datasets/) with --ortholog flag. Retrieval of data was automated by bash script. All retrieved genes were checked if the retrieved gene symbol is identical to the query. In the case of any discrepancy between searched and retrieved gene symbols, their equivalency was further investigated. All genes missing in ortholog databases in chicken, in paleognath birds or in Crocodylia were searched by tBLASTn in the genomes of chicken huxu breed (GCA_024206055.2), ostrich (GCA_040807025.1), and american alligator (GCA_030867095.1), respectively, using protein query from an evolutionary close vertebrate species. If any missing gene was found, the quality of the match and syntenic context was used to assess the validity of the gene found. Any good match in appropriate syntenic context was used as proof of the gene presence without verification if the gene is functional or if it is pseudogene.

### Repeat content analysis

Repeat annotation data were downloaded from UCSC GB hub directory (Clawson et al. 2023)(https://hgdownload.soe.ucsc.edu/hubs/GCA/024/206/055/GCA_024206055.2).

Combined simple repeat annotation was created with a custom script selecting ranges of ‘Simple_repeats’ class from the RepeatMasker annotation (GCA_024206055.2.repeatMasker.out.gz, version 2024-12-03) and the resulting annotation was merged with Tandem Repeat Finder (Benson 1999) annotation (GCA_024206055.2_GGswu.simpleRepeat.bb, version 2024-12-03). Intron coverage by simple repeats was calculated using pybedtools coverage command with default parameters using intron annotation (see Exon stuttering screen) and combined simple repeat annotation as inputs.

### BLASTn dot plots

For BLASTn dotplots, genomic sequences were compared to itself or to virtually spliced cDNA sequence based on the gene annotation. The source of the sequences were either reference genome assembly or reconstructed sequences from nanopore sequencing reads as specified in the figure legends. The BLASTn search algorithm (Camacho et al. 2009) was performed with low complexity regions filter switched off and other parameters set to default values. To determine changes in sequence structure for individual intron alleles of chicken genes we manually aligned allele sequences based on the repeat clusters identified in the BLASTn dotplots. This solution is only approximate and was not intended to reveal variation on the level of individual base pairs.

### Data processing and visualization

Results of exon stuttering screen data (partial exon copies in adjacent introns of chicken genes) were stored in the interact file format as specified by UCSC Genome Browser (Perez et al. 2025) and visualised as an annotation track. Data were processed using R version 4.5.1 (R Core team 2025). Data were processed through custom MySQL scripts using DBI (Wickham et al. 2024) and RMariaDB (Müller et al. 2017) packages. Visualisations (box plots and bar plots) were created using ggplot2 (Wickham 2009) package.

### Exon stuttering screen

Exon and intron annotations were created from XenoRef annotation GTF file (see Reference genomes and gene annotations) with custom script. For each gene, marked by gene ID, exon entries are identified, grouped by gene ID, and stored in a BED file. To define introns, the exons for each gene were sorted by their genomic start positions. The ranges of length ≥ 10 between consecutive exons were considered as introns and were saved as BED files. Exon and intron sequences were extracted from the GCA_024206055.2 reference genome sequence using pybedtools sequence command with s=True option. For the exon stuttering screen, only exons with no overlap with repetitive elements (including all but ‘Simple_repeat’ and ‘Low_complexity’ ranges from the RepeatMasker annotation, see Repeat content analysis for annotation source) were included. As a control for unspecific hit characteristics, shuffled intron sequences (the shuffled control) were generated from intron sequences using the function shuffle from a python module random.

Exon stuttering screen was performed using a BLASTn-based approach. First, for each gene, exonic and intronic sequences were extracted from their respective FASTA files. BLASTn (version 2.12.0) was then run with -task blastn-short -dust “no” options, where each exon sequence was used as a query against a set of all introns of the respective gene as a subject. Only hits with length ≥ 20 were accepted. Identically, each exon was compared to a set of shuffled intron sequences. Hits were converted to BED-like format with genomic coordinates. Intronic hits (targets) with more than one source range (exon ranges) were discarded.

### Intron length analysis from nanopore sequencing data

The following publicly available human datasets from the NCBI Short Read Archive (SRA) were used (SRA study PRJNA1108179): SRR30344988, SRR30344989, and SRR30344990. The following publicly available chicken datasets from the NCBI SRA were used (chicken breed and SRA study in parentheses): SRX11722867 (Huxu, PRJNA693184), SRX11722868 (Huxu, PRJNA693184), SRX11722871 (Huxu, PRJNA693184), SRX14125033 (Silkie, PRJNA805080), SRX14125034 (Silkie, PRJNA805080), SRX19311940 (ACRB, PRJNA694114), SRX19311957 (Cornell2, PRJNA694114), SRX19311958 (Luoyangwu, PRJNA694114), SRX19311959 (Araucana, PRJNA694114), SRX19311960 (red junglefowl, PRJNA694114), and SRX19311961 (Tibetan, PRJNA694114). All datasets were generated by nanopore technology and of sufficient size to allow large scale analysis of introns.

Sequencing data for individual samples were converted to fasta format and mapped to the corresponding genome reference using minimap2 v. 2.26-r1175 (H. Li 2018) with following parameters: --secondary=no -a -t 18 -x map-ont. Based on the corresponding gene annotation, a list of regions comprising introns and their adjacent exons was created. Introns with length <10bp were excluded. Exons of overlapping genes were merged and treated as single gene exons. Reads for each region were retrieved and positions of each exon were identified by BLASTn search of reference exon sequences with following criterias: BLASTn bit score > 50, ii) hit alignment cannot differ more than 20% from reference exon length.

Length of intron region (region between exon positions identified) was calculated for each sequencing read where both exons were identified in the same orientation. Lists of lengths for each intron were then clustered into the 2 categories (2 alleles) by Jenks optimization method (Jenks 1967). Length of individual alleles were defined as means of lengths in each category.

As a control, artificial sequencing data for human and chicken were simulated from the reference genome sequences by NanoSim v. 3.2.2 (Yang et al. 2017) with sequencing coverage 100x and analyzed in parallel to the real samples. These datasets should not contain any real variability because they are produced from the haploid references. All detected variability was therefore considered as an artifact of the method (mainly caused by false identification of exons of sequentially similar genes).

To reduce false positives, each intron in each sample had to pass the following criteria: i) intron length > 100b for at least one allele, ii) >4 reads identified for the intron, iii) read counts for individual alleles cannot differ more than 10 times, iv) all intron lengths identified for each allele cannot differ more than 10% from mean allele length, and v) all introns detected in the simulated datasets were excluded. These criteria were determined empirically to maximize the screen specificity. After filtering, we obtained informative results for 70% of all annotated chicken introns and for 49% of human introns.

As a result, length alleles for individual introns in individual samples were obtained. In the final step, length alleles for each intron were aggregated across the samples to produce sets of different alleles in the sampled population. Unique allele was defined as an allele with intron length that differs from any other allele by more than 10%. This criterion was used to exclude false-positive alleles produced by sequencing indel errors. Allele richness (number of alleles) for each intron was then rarefied to n=6 (3 diploid samples) and used for identification of population variability of intron lengths.

