## Supplementary figures for "Decoding the avian missing gene mystery: dot chromosomes unmask extensive gene loss and novel genetic instability"

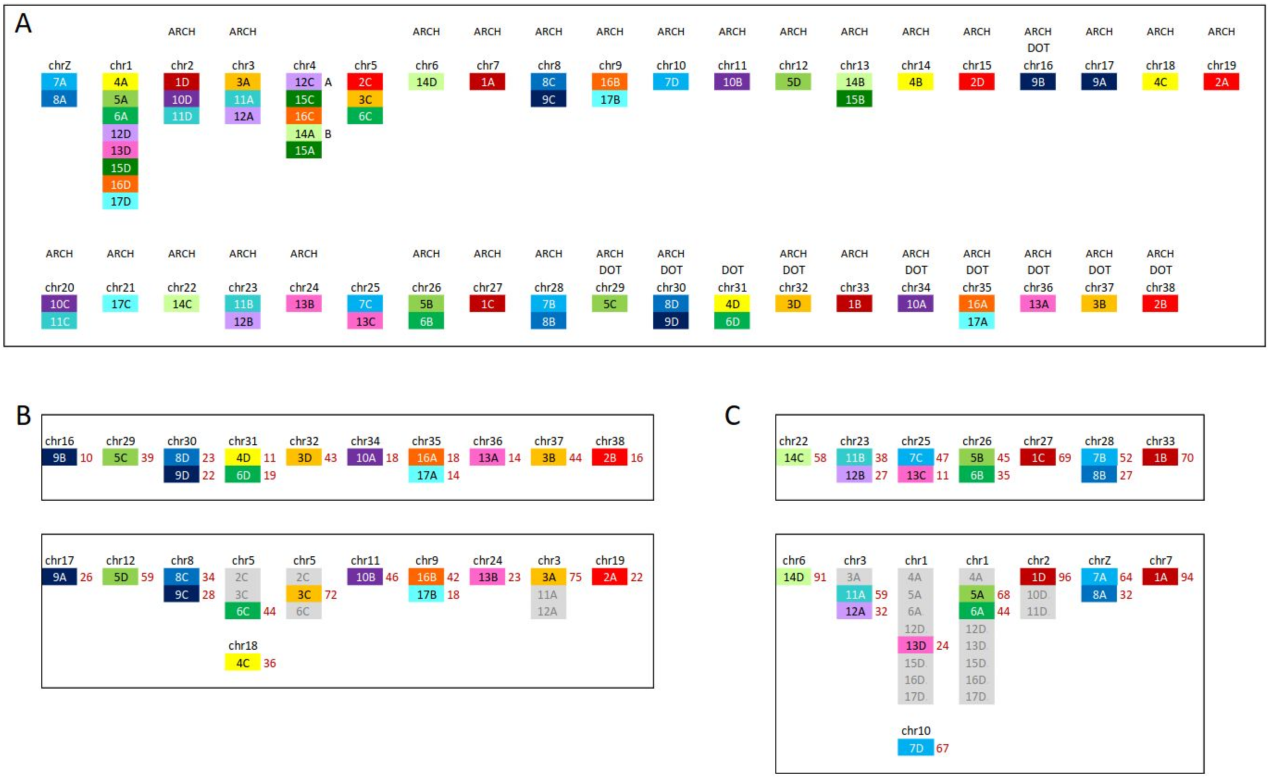


**Supplementary Figure 1. Paralogon maps.** System of 68 paralogons (17 prevertebrate paralogons, each quadrupled) as defined by Lamb 2021 (1). (A) Paralogon map of chicken genome. Thirty nine chicken chromosomes (2) composed of 38 autosomes and sex chromosome Z (chrW omitted) with corresponding paralogons shown as color-coded rectangles. Dot chromosomes (DOT) and archeochromosomes (ARCH) are denoted above specific chromosomes. Archeochromosomes as defined by Lamb 2021 are chromosomes that did not underwent any interchromosomal fusion or fission from the time of the last common ancestor of contemporary jawed vertebrates. Chicken chr4 is formed by fusion of two chromosomes (A:12C-15C-16C, B:14A-15A) that were originally separate and as separate chromosomes exist in many avian species. (B) Paralogon map of 10 chicken dot chromosomes with chromosomes carrying corresponding isoparalogons shown below. On the right side of each rectangle representing one paralogon is a number of ohnologs used for analysis presented in Fig. 1 and Suppl. Tab. 1 and Suppl. Tab. 2. (C) Paralogon map of non-Dot chromosomes smaller than 7Mb with chromosomes carrying corresponding isoparalogons shown below. On the right side of each rectangle representing one paralogon is a number of ohnologs used for analysis presented in Suppl. Fig. 3

1. Lamb, Trevor D. 2021. "Analysis of Paralogons, Origin of the Vertebrate Karyotype, and Ancient Chromosomes Retained in Extant Species." Genome Biology and Evolution 13 (4). https://doi.org/10.1093/gbe/evab044.
2. Huang, Zhen, Zaoxu Xu, Hao Bai, Yongji Huang, Na Kang, Xiaoting Ding, Jing Liu, et al. 2023. "Evolutionary Analysis of a Complete Chicken Genome." Proceedings of the National Academy of Sciences of the United States of America 120 (8): e2216641120.

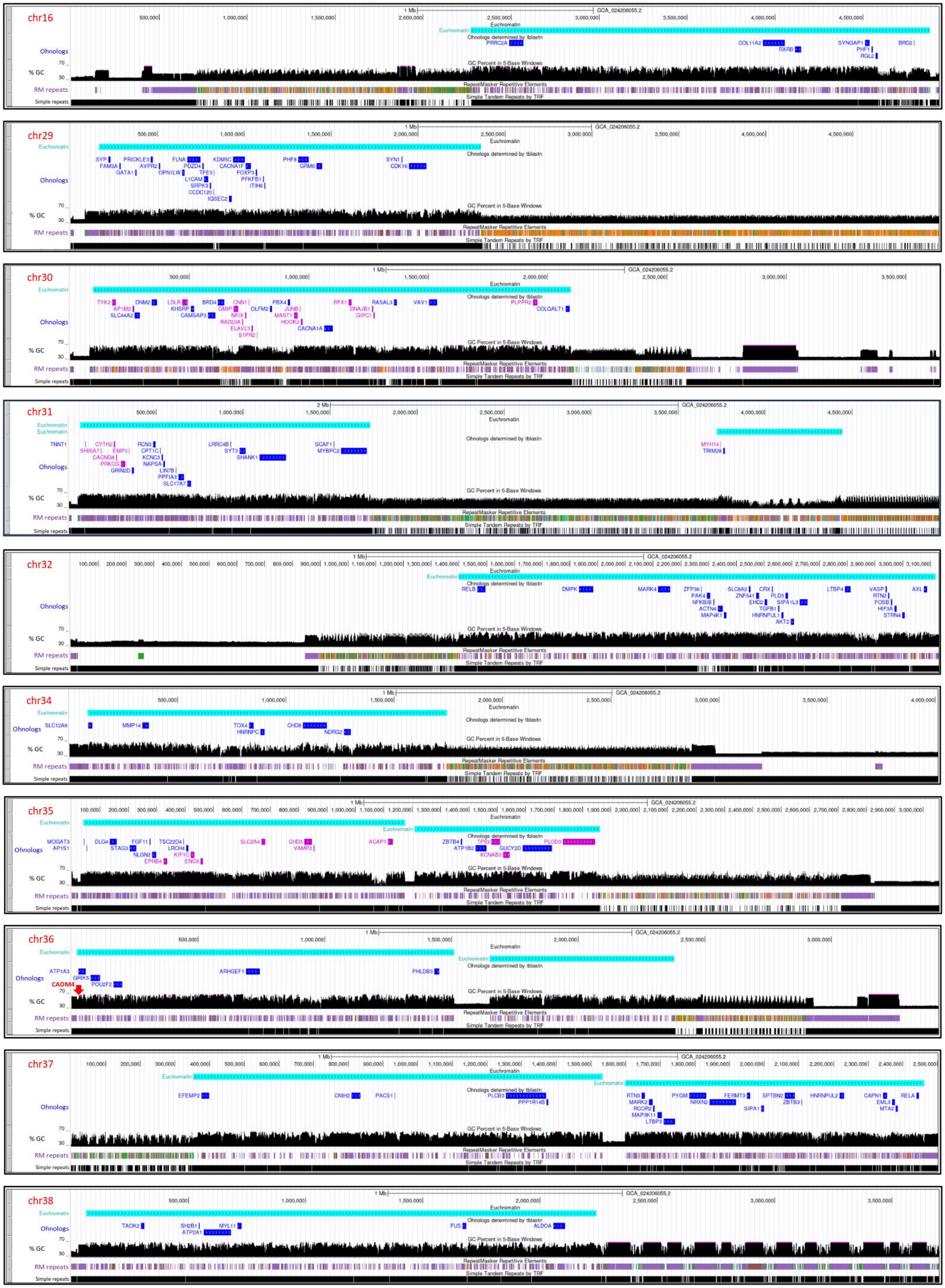


**Supplementary Figure 2.** See the legend on the next page.

**Supplementary Figure 2. Location of ohnologs found on chicken dot chromosomes relative to positions of euchromatin regions.** The figure shows screenshots of chicken huxu dot chromosomes visualized in UCSC genome browser (https://genome.ucsc.edu/). If the dot chromosome was longer than 5 Mb (chr29, chr31), only first 5Mb of the sequence are shown. Tracks showing % of GC content, repeats detected by Repeat Masker (RM repeats) and repeats detected by Tandem Repeat Finder are internal tracks provided by UCSC genome browser. BED file of ohnologs shows coding sequences including intervening introns. The positions of these regions were determined by blasting protein sequences of avian orthologs identified by NCBI ortholog database against GGswu genome using tblastn. Regions are shown in blue ink if ohnologs came from the same paralogon. If there are ohnologs from two paralogons on the same chromosome blue and violet inks are used to differentiate ohnolog‘s origin. Short region of genomic sequence that contains CADM4 gene is missing from chromosome 36 in huxu genome. The position of missing DNA stretch is indicated by red arrow. Positions of putative euchromatin are shown by cyan blue rectangles positioned based on the published schema of dot chromosome compartmentalization (1). The positioning was further precised using specific repeat content of dot euchromatin regions (high simple tandem repeats excluding long subtelomere stretches, low content of interspersed repeats).

1. Huang, Zhen, Zaoxu Xu, Hao Bai, Yongji Huang, Na Kang, Xiaoting Ding, Jing Liu, et al. 2023. “Evolutionary Analysis of a Complete Chicken Genome.” Proceedings of the National Academy of Sciences of the United States of America 120 (8): e2216641120.

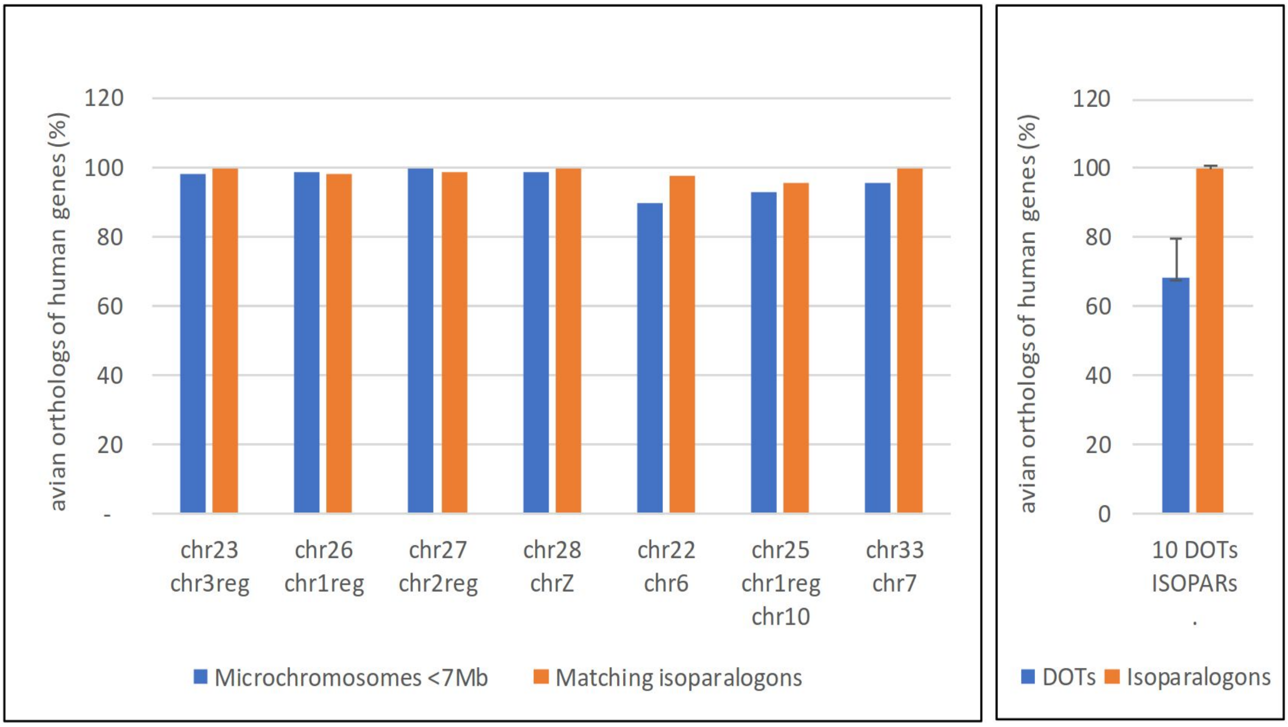


**Supplementary Figure 3. Survival of ohnologs in avian paralogons mapping to chicken chromosomes smaller than 7Mb.** Survival in matching isoparalogons on other chromosomes is shown as well. The figure compares survival in paralogons mapping to individual non-dot chromosomes and corresponding isoparalogons with surviving ohnologs averaged through 10 dot chromosomes (10 DOTs) and matching isoparalogons on other chromosomes (ISOPARs). Error bars represent standard deviation. Designations of individual small non-dot chromosomes are indicated as well as designations of chromosomes with matching isoparalogons shown below. If isoparalogons don‘t cover entire chromosome suffix reg (region) is added, e.g. chr3reg. semiDOTs is a special category of non-dot microchromosomes which have a set of features shared with dot chromosomes.

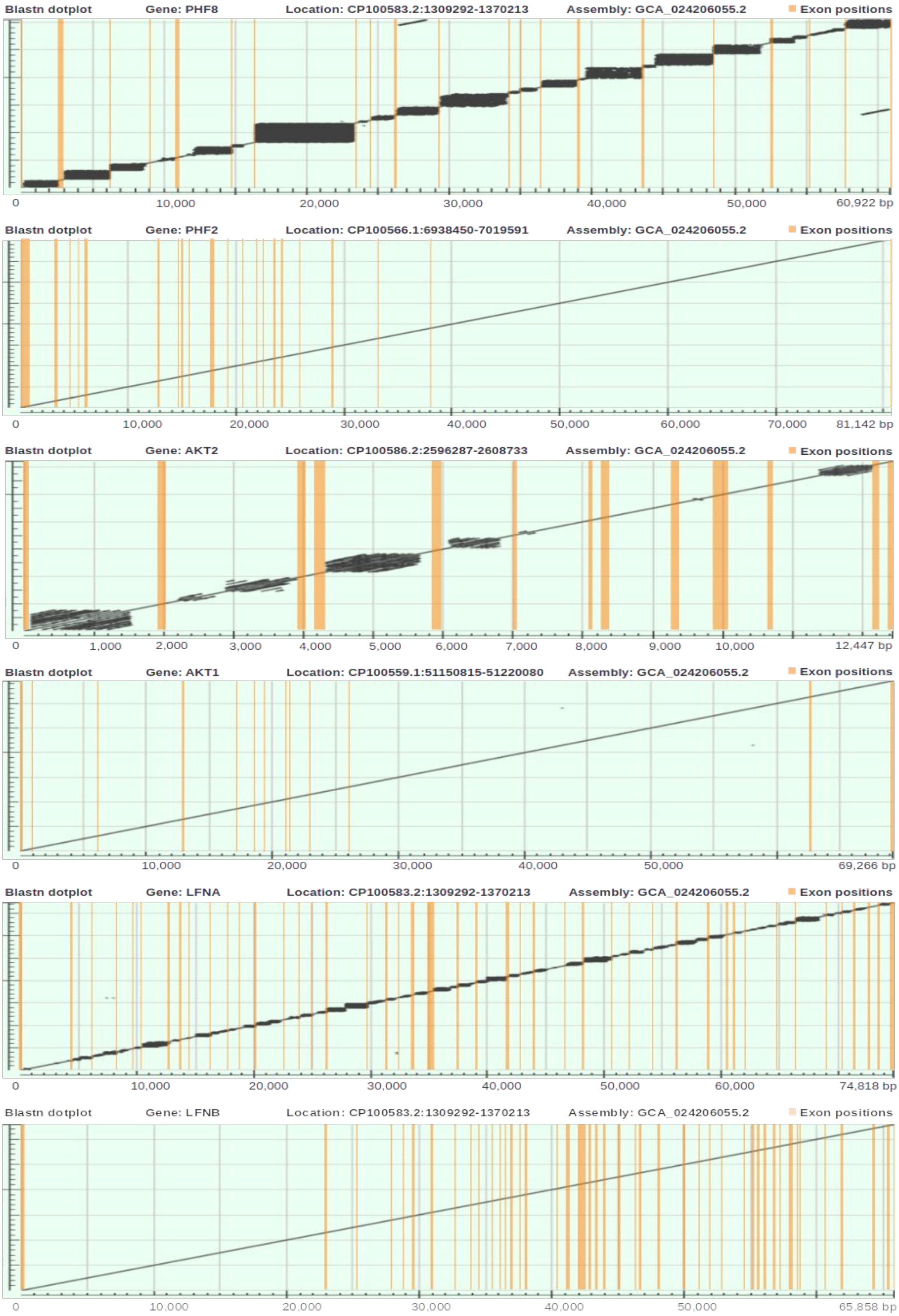


**Supplementary Figure 4. Example of local repeat expansion in chicken genes PHF8, AKT2 and LFNA located on dot chromosomes, and their ohnologs PHF2, AKT1 and LFNB located on other chromosomes.** Sequence dot plots are shown for blastn search of genomic sequence vs. itself (upper schemes), and cDNA vs. genomic sequence (lower schemes). Dark regions in the plots represent blastn hits. Exon positions are manually curated and indicated by an orange background.

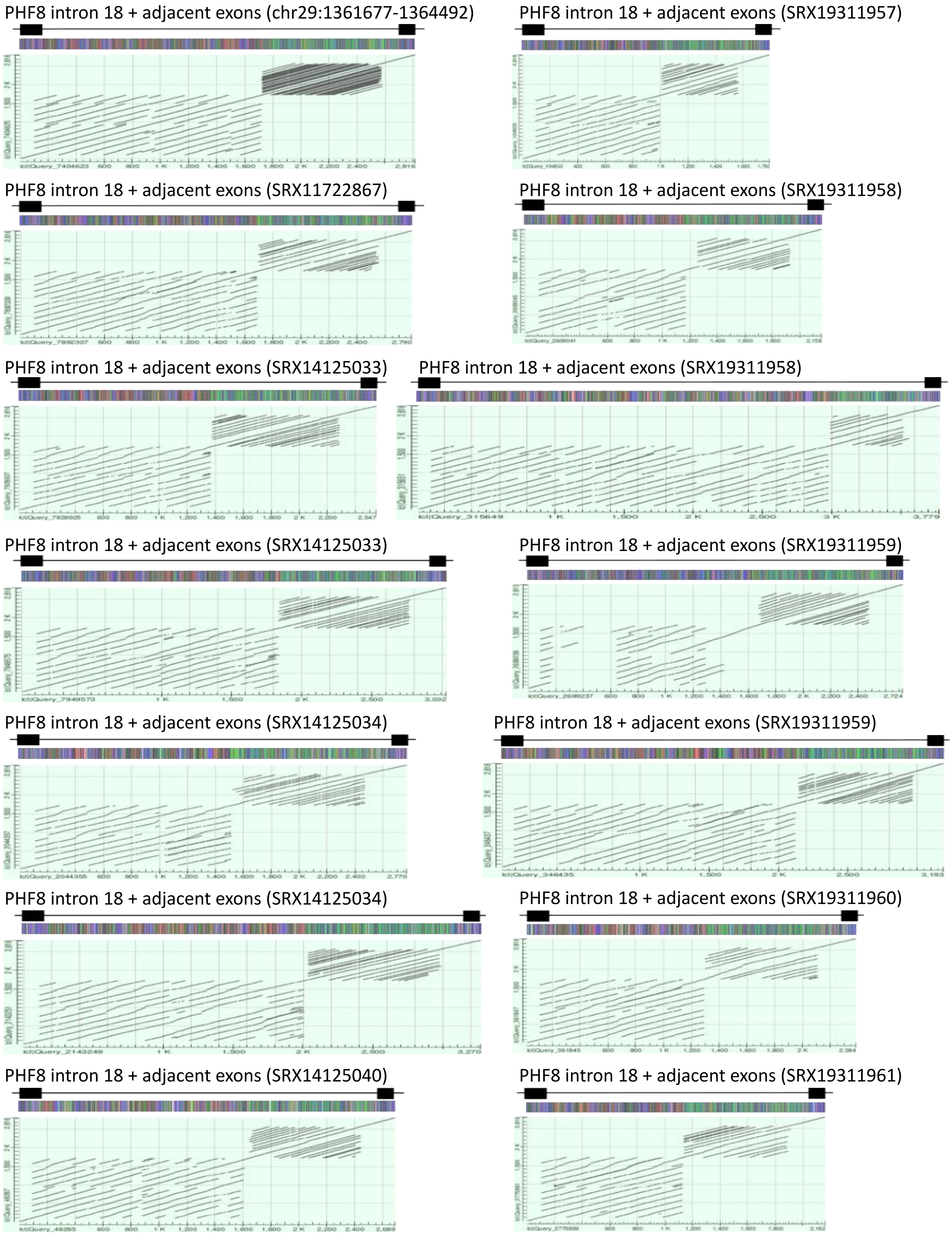


**Supplementary Figure 5. Sequence structure of individual alleles of chicken PHF8 intron 18.** Sequence dot plots are shown for blastn self-alignments of allele sequences comprising intron and adjacent exons. First chart represents Ggswu reference sequence. Allele sequences were reconstructed from Nanopore sequencing of chicken individuals. Allele sequences are shown in a colored schemes above each chart where each nucleotide is represented by a different color (A - green, C - blue, G - grey, T - red).

| Paralogons | Spotted gar | Reedfish | Xenopus | Human | Spotted gar | Reedfish | Xenopus | Human | Spotted gar | Reedfish | Xenopus | Human | Spotted gar | Reedfish | Xenopus | Human |
|---|---|---|---|---|---|---|---|---|---|---|---|---|---|---|---|---|
| | A | A | A | A | B | B | B | B | C | C | C | C | D | D | D | D |
| 1 | 96 | 85 | 84 | 94 | 73 | 67 | 67 | 70 | 70 | 69 | 57 | 70 | 96 | 96 | 88 | 96 |
| 2 | 23 | 22 | 23 | 22 | 6 | 15 | 8 | 16 | 11 | 10 | 2 | 10 | 21 | 20 | 22 | 23 |
| 3 | 75 | 70 | 67 | 75 | 34 | 42 | 33 | 44 | 71 | 71 | 69 | 73 | 46 | 39 | 32 | 44 |
| 4 | 20 | 28 | 25 | 30 | 35 | 31 | 34 | 33 | 37 | 34 | 33 | 37 | 6 | 13 | 13 | 11 |
| 5 | 66 | 64 | 65 | 68 | 47 | 43 | 39 | 45 | 35 | 40 | 34 | 39 | 60 | 56 | 55 | 59 |
| 6 | 48 | 47 | 39 | 44 | 39 | 39 | 25 | 36 | 45 | 46 | 43 | 44 | 4 | 14 | 18 | 19 |
| 7 | 69 | 66 | 62 | 64 | 57 | 53 | 48 | 52 | 43 | 45 | 40 | 47 | 70 | 59 | 62 | 67 |
| 8 | 33 | 30 | 33 | 32 | 30 | 27 | 23 | 27 | 35 | 35 | 34 | 35 | 26 | 23 | 21 | 23 |
| 9 | 26 | 22 | 25 | 26 | 1 | 6 | 7 | 10 | 27 | 25 | 24 | 28 | 22 | 20 | 17 | 22 |
| 10 | 5 | 12 | 11 | 19 | 50 | 49 | 44 | 47 | 45 | 42 | 34 | 39 | 51 | 46 | 45 | 50 |
| 11 | 60 | 59 | 52 | 60 | 43 | 40 | 38 | 38 | 48 | 48 | 30 | 46 | 63 | 54 | 60 | 62 |
| 12 | 32 | 32 | 29 | 32 | 26 | 25 | 23 | 27 | 25 | 24 | 24 | 19 | 34 | 34 | 33 | 34 |
| 13 | 16 | 14 | 15 | 14 | 23 | 22 | 23 | 23 | 6 | 6 | 8 | 11 | 25 | 20 | 16 | 24 |
| 14 | 94 | 89 | 86 | 88 | 72 | 69 | 57 | 68 | 61 | 58 | 48 | 58 | 93 | 87 | 82 | 91 |
| 15 | 53 | 52 | 49 | 56 | 42 | 42 | 40 | 38 | 41 | 38 | 37 | 31 | 55 | 51 | 50 | 56 |
| 16 | 21 | 18 | 11 | 18 | 43 | 40 | 38 | 42 | 40 | 34 | 40 | 41 | 44 | 42 | 40 | 42 |
| 17 | 14 | 14 | 8 | 14 | 18 | 18 | 19 | 19 | 21 | 20 | 17 | 22 | 12 | 11 | 10 | 14 |

| Summary (average) for all four species | | | | |
|---|---|---|---|---|
| Paralogons | A | B | C | D |
| 1 | 89.8 | 69.3 | 66.5 | 94 |
| 2 | 22.5 | 11.3 | 8.25 | 21.5 |
| 3 | 71.8 | 38.3 | 71 | 40.3 |
| 4 | 25.8 | 33.3 | 35.3 | 10.8 |
| 5 | 65.8 | 43.5 | 37 | 57.5 |
| 6 | 44.5 | 34.8 | 44.5 | 13.8 |
| 7 | 65.3 | 52.5 | 43.8 | 64.5 |
| 8 | 32 | 26.8 | 34.8 | 23.3 |
| 9 | 24.8 | 6 | 26 | 20.3 |
| 10 | 11.8 | 47.5 | 40 | 48 |
| 11 | 57.8 | 39.8 | 43 | 59.8 |
| 12 | 31.3 | 25.3 | 23 | 33.8 |
| 13 | 14.8 | 22.8 | 7.75 | 21.3 |
| 14 | 89.3 | 66.5 | 56.3 | 88.3 |
| 15 | 52.5 | 40.5 | 36.8 | 53 |
| 16 | 17 | 40.8 | 38.8 | 42 |
| 17 | 12.5 | 18.5 | 20 | 11.8 |

| Chromosomal location of paralogons | | | | |
|---|---|---|---|---|
| Paralogons | A | B | C | D |
| 1 | 7 | 33 | 27 | 2 |
| 2 | 19 | 38 | 5 | 15 |
| 3 | 3 | 37 | 5 | 32 |
| 4 | 1 | 14 | 18 | 31 |
| 5 | 1 | 26 | 29 | 12 |
| 6 | 1 | 26 | 5 | 31 |
| 7 | Z | 28 | 25 | 10 |
| 8 | Z | 28 | 8 | 30 |
| 9 | 17 | 16 | 8 | 30 |
| 10 | 34 | 11 | 20 | 2 |
| 11 | 3 | 23 | 20 | 2 |
| 12 | 3 | 23 | 4 | 1 |
| 13 | 36 | 24 | 25 | 1 |
| 14 | 4 | 13 | 22 | 6 |
| 15 | 4 | 13 | 4 | 1 |
| 16 | 35 | 9 | 4 | 1 |
| 17 | 35 | 9 | 21 | 1 |

**Supplementary Figure 6. Biased fractionation of numbers of identified ohnologs suggest that chicken dot chromosomes come from recessive subgenome of the second WGD karyotype.** The table on the left shows number of ohnologs found for each paralogon and quad in four non-archosaurian species. Data comes from the Supplementary table 1B published by (1). The second table averages the numbers through all four species. Because quad pairs A-B and C-D were generated to reflect the second WGD, one of the quads of A-B pair or C-D pair comes from dominant genome while the other of the same pair from recessive genome. Biased fractionation indicates that the quad containing higher number of ohnologs (on green background) comes from dominant while the quad with lower number of ohnologs (on reddish brown background) comes from recessive subgenome. The table on the right shows chromosomal location of each paralogon/quad in chicken. Dot chromosomes are indicated by red ink on yellow background.

1. Lamb, Trevor D. 2021. "Analysis of Paralogons, Origin of the Vertebrate Karyotype, and Ancient Chromosomes Retained in Extant Species." Genome Biology and Evolution 13 (4). https://doi.org/10.1093/gbe/evab044.
